# Gene mobility elements mediate cell type specific genome organization and radial gene movement *in vivo*

**DOI:** 10.1101/2024.11.30.626181

**Authors:** Tanguy Lucas, Lin-Ing Wang, Juniper Glass-Klaiber, Elvis Quiroz, Sofiya Patra, Natalia Molotkova, Minoree Kohwi

## Abstract

Understanding the level of genome organization that governs gene regulation remains a challenge despite advancements in chromatin profiling techniques. Cell type specific chromatin architectures may be obscured by averaging heterogeneous cell populations. Here we took a reductionist perspective, starting with the relocation of the *hunchback* gene to the nuclear lamina in *Drosophila* neuroblasts. We previously found that this event terminates competence to produce early-born neurons and is mediated by an intronic 250 base-pair element, which we term gene mobility element (GME). Here we found over 800 putative GMEs globally that are chromatin accessible and are Polycomb (PcG) target sites. GMEs appear to be distinct from PcG response elements, however, which are largely chromatin inaccessible in neuroblasts. Performing *in situ* Hi-C of purified neuroblasts, we found that GMEs form megabase-scale chromatin interactions, spanning multiple topologically associated domain borders, preferentially contacting other GMEs. These interactions are cell type and stage-specific. Notably, GMEs undergo developmentally- timed mobilization to/from the neuroblast nuclear lamina, and domain swapping a GFP reporter transgene intron with a GME relocates the transgene to the nuclear lamina in embryos. We propose that GMEs constitute a genome organizational framework and mediate gene-to-lamina mobilization during progenitor competence state transitions *in vivo*.

## INTRODUCTION

The metazoan genome is non-randomly organized, and it is thought that its three dimensional folding and packaging underlie cell type specific gene expression program. Advancements in chromatin profiling approaches have uncovered the key principles of genome organization at multiple scales. Yet, what levels of genome organization relate to gene regulation and cell functions remain largely unknown, as biologically relevant chromatin interactions may be rare and difficult to identify without cell and stage-specific experimentation (Finn and Misteli, 2019; Finn et al., 2019; Kohwi et al., 2013; Lucas et al., 2021; Lucas and Kohwi, 2019; Vermunt et al., 2018). In addition to understanding its role in distal enhancer-promoter interactions underlying transcriptional activity (Cavalheiro et al., 2021; Levo et al., 2022), genome organization can also mediate epigenetic changes that alter whether a gene is poised to be activated upon changes in cell states, like differentiation (Meister et al., 2010) or changes in states of competence that affect gene expression of future descendent cells (Kohwi et al., 2013; Lucas et al., 2021). Thus, how and what level of genome organization is linked to different types of gene regulation is a complex question. It is a particularly challenging problem for neural progenitors, which generate diverse descendent cell types with distinct gene expression profiles over the course of their lineage.

Progress in identifying higher-order principles of genome folding has focused on topologically associated domains (TADs), which are thought to establish subregions of the genome with higher interaction frequencies, in order to both facilitate and insulate chromatin contacts (Dixon et al., 2012; Hou et al., 2012; Nora et al., 2012; Sexton et al., 2012). However, recent studies reveal that TADs may instead be an emergent statistical property from averaging populations of cells whose chromatin conformations are highly dynamic, rather than a stable structure (Bintu et al., 2018; Chen et al., 2023; Finn and Misteli, 2019). Further, TADs can be conserved across cell types (Dekker and Heard, 2015; McArthur and Capra, 2021; Rao et al., 2014; Schmitt et al., 2016), raising the possibility that additional genome organization features might play a role in facilitating cell type specific gene regulation. In addition to TADs, lamina- associated domains (LADs) have also emerged as a type of chromatin domain, in which LADs are defined by interaction with the nuclear lamina and genes within LADs have been found to be highly associated with silenced or repressed states (Alagna et al., 2023; Gonzalez-Sandoval and Gasser, 2016; Harr et al., 2015; Pickersgill et al., 2006; van Steensel and Belmont, 2017). While TADs and LADs have been identified as common genome organizational units of animal genomes, an *in vivo* approach that considers cell function and developmental gene regulation is still necessary to provide context specificity to the relationship between genome architecture and gene function.

In *Drosophila*, embryonic neuroblasts (neural progenitors) sequentially produce distinct neural cell types by expressing, as they divide, a series of transcriptional factors, called temporal identity factors (Doe, 2017; Isshiki et al., 2001; Kohwi and Doe, 2013). In multiple neuroblast lineages, Hunchback (Hb) is the first of the temporal identity factors to be expressed, and it specifies early-born identity of the neuronal progeny (Brody and Odenwald, 2000; Grosskortenhaus et al., 2005; Isshiki et al., 2001; Kohwi and Doe, 2013; Kohwi et al., 2013). Hb acts in the neuroblast to “prime” the future transcriptional program of the descendent neuron, which includes transcribing the *hb* gene, a “molecular time stamp” of neuron birth order (Grosskortenhaus et al., 2005; Kohwi et al., 2013). Over time, neuroblasts transit through distinct states of competence, which limits the developmental time period in which each cell type can be specified (Cleary and Doe, 2006; Kohwi et al., 2013; Pearson and Doe, 2003). We previously discovered that loss of neuroblast competence to produce early-born neurons was due to the developmentally-timed relocation of the *hb* gene to the neuroblast nuclear periphery, which occurs several divisions and hours after the *hb* gene is already transcriptionally repressed in the neuroblast. Instead, the relocation event results in the heritable silencing of the *hb* gene where it becomes refractory to activation in the descendent neurons born two mitotic divisions later (Hafer et al., 2022; Kohwi et al., 2013; Lucas et al., 2021). Thus, changes in organization of the progenitor genome alter its potential to regulate gene expression programs in the descendent progeny.

We recently identified a 250bp *cis*-acting element in the *hb* intron that is necessary and sufficient for *hb* gene relocation to the nuclear lamina (Lucas et al., 2021), which we call <u>G</u>ene <u>M</u>obility <u>E</u>lement (GME). Here, informed by the characteristics of *hb* GME, we identify putative GMEs genome-wide, and explore how they are organized within the neuroblast genome *in viv*o. Using *in situ* high throughput chromatin conformation capture (Hi-C) on neuroblasts purified from embryos, we found that GMEs are largely associated with neuronal genes, and strongly interact across tens of megabases of distances, traversing multiple topologically associated domain (TAD) borders. Further, we observed that GME interactions are cell type specific and are dynamic in neuroblasts across time. Using targeted Lamin DamID (Southall et al., 2013; van Steensel and Henikoff, 2000) we found that GMEs change their association with the neuroblast nuclear lamina in late-stage embryos. Finally, we tested two putative GMEs and found that they are both individually sufficient to promote the relocation of a reporter transgene to the nuclear lamina and to repress reporter expression *in vivo,* highly consistent with the function described for the *hb* GME in the embryo during competence closure. Together, the results reveal a framework for cell type and stage-specific genome organization that mediates relocation of genes to the nuclear lamina *in vivo*.

## RESULTS

### Genomic elements similar to *hb* GME are found genome-wide

In neuroblasts, the mid-embryonic relocation of the *hb* gene to the nuclear periphery causes its heritable silencing and terminates competence to generate descendent neurons that can transcribe *hb* (Kohwi et al., 2013; Lucas et al., 2021) **(Fig. 1A)**. We previously identified a 250bp element in the *hb* intron that mediates *hb* gene relocation to the nuclear lamina and that requires Polycomb (PcG) function. PcG factors are key players of epigenetic processes and are known to exert their well-studied roles in maintenance of gene repression via the histone mark H3 lysine 27 trimethylation modification (H3K27me3) (Grossniklaus and Paro, 2014). In neuroblasts, PcG factors of both PcG repressive complex 1 and 2 (PRC1 and PRC2) are expressed, and yet neuroblast nuclei are markedly devoid of H3K27me3 signals (Abdusselamoglu et al., 2019; Arya et al., 2019; Lucas et al., 2021). Immunostaining for this mark in embryos shows that neuroblasts have fewer and weaker H3K27me3 puncta quantified compared to non-neuroblast cells (**Supp Fig. 1A**). Moreover, established Hox target genes such as Abd-B are not de-repressed in severe PRC2 mutant embryo neuroblasts compared to the neighboring epithelial cells (**Supp Fig. 1B**). These observations are consistent with our previous report that loss of PRC1 did not affect *hb* transcriptional activity, only its relocation to the nuclear lamina, and suggest a genome architecture function of PcG factors in neuroblasts. We hypothesized that perhaps there exists a group of genomic elements that, like the *hb* GME, may be a target of PcG chromatin factors but act in a genome organizational role. We thus sought to identify such sequences using the characteristics of the *hb* intronic GME as reference. We selected sequences that showed high chromatin accessibility in neuroblasts and being a Polycomb (PcG) chromatin factor target site. We used published ChIP-seq datasets from the larval brain (Brown et al., 2018) to identify significant binding targets for the core PRC1 subunit, Psc, core PRC2 subunit, Enhancer of zeste, E(z), and Pho, the only known PcG subunit that directly binds chromatin (Brown et al., 1998). Additionally, a recent study showed that the nuclear pore complex subunit Nup93 interacts with PcG factors to silence target genes at the nuclear periphery (Pascual-Garcia et al., 2017). Using the larval CNS Nup93 ChIP-seq dataset (Pascual-Garcia et al., 2017), we found that the *hb* intronic element (*hb* GME) is also a Nup93-target site (**Fig. 1B**). Based on these characteristics, we identified genome wide 859 sites highly similar to the *hb* GME, which we will refer collectively to hereafter as “GME” (**Supp Data 2**). PcG factors are best known for their role in the maintenance of gene silencing by binding to PcG response elements (PREs) (Chan et al., 1994; Grossniklaus and Paro, 2014; Simon et al., 1993). A recent study using machine-learning based algorithms have identified PREs in the fly genome by incorporating experimental and bioinformatic analyses of multiple published datasets (Bredesen and Rehmsmeier, 2019). We compared their list of the core 2908 PREs with our list of 859 GMEs and found the two are generally non-overlapping. Notably, PREs were largely in a chromatin inaccessible state in neuroblasts (**Fig. 1C**). Additionally, we found that the majority of genes (70%) that were in close proximity to GMEs were characterized as neuronal genes, and 53% were associated with embryonic development, nearly twice as many as expected for neurodevelopmental genes in the whole genome (**Fig. 1D, Supp Data 1 and 3**).

**Figure 1:**
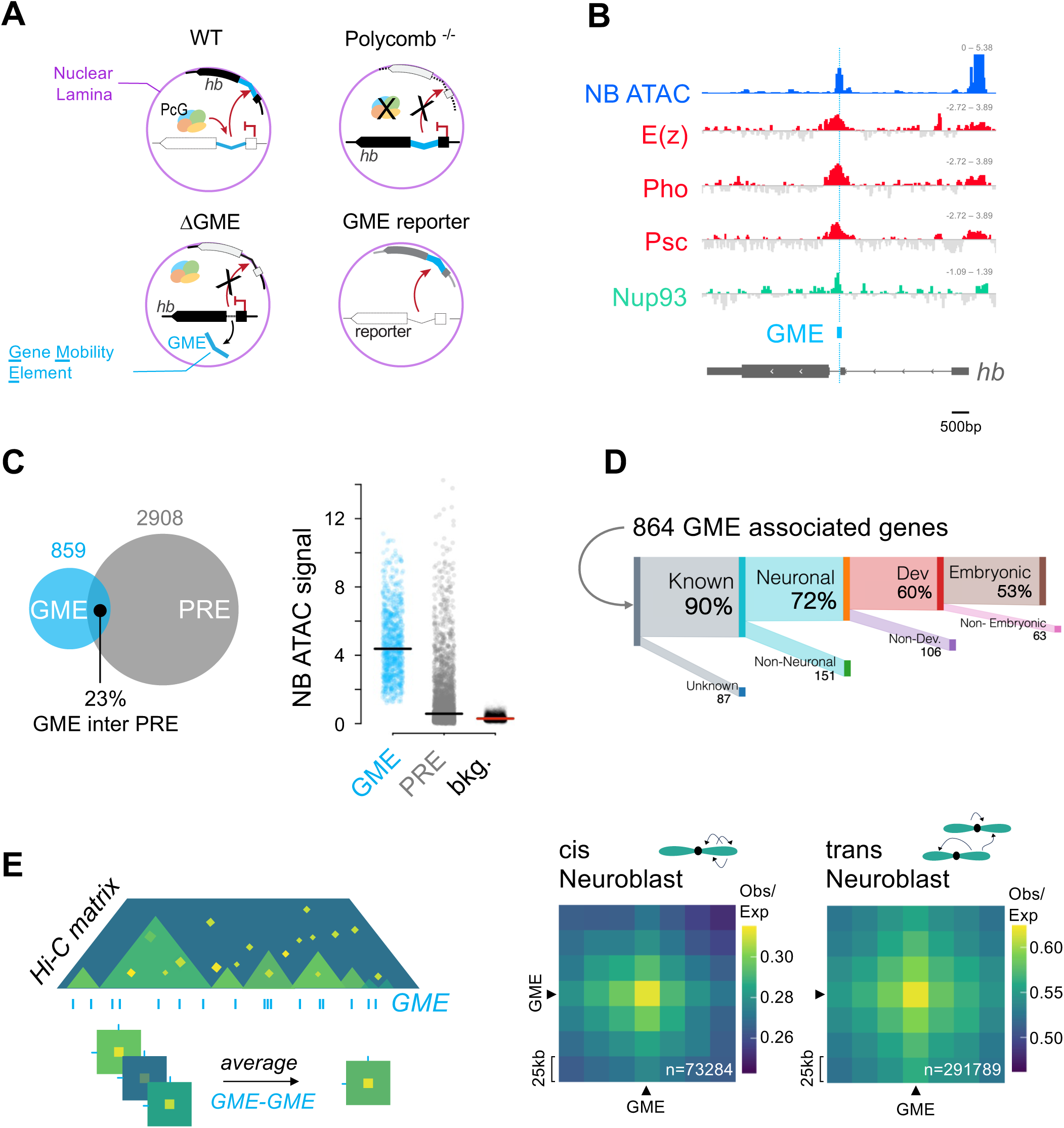
Identification of putative GME genome wide. A. Schematic adapted from Lucas et al., 2021. The *hunchback* (*hb*) GME, cyan, located in the (*hb*) intron, is both necessary (ΔGME) and sufficient (GME reporter context) for the relocation of the *hb* gene to the nuclear lamina (magenta) in neuroblasts. The Polycomb repressive complex 1 subunit Psc is required for *hb* relocation to the nuclear lamina. **B. Polycomb Factors are enriched at the *hb* GME:** Genomic tracks at the *hb* locus centered on the *hb* GME (cyan): Chromatin accessibility profile by ATACseq in embryonic stage 14 neuroblasts in blue; ChIP-seq profiles from L3 Larval CNS (Brown et al., 2018) of E(z) (PRC2 core subunits), Pho and Psc (PRC1 core subunit) in red; and nuclear pore complex protein Nup93 (Pascual-Garcia et al., 2017) in green. **C. GMEs and Polycomb Response Elements (PREs) are distinct:** *left*: Venn diagram showing the overlap between GMEs identified genome-wide and PREs (Bredesen *et al*., 2018) (overlap: n = 202 of 859 total GMEs, 23%). *Right*: Neuroblast ATACseq signal (averaged from embryonic stage 10, 12 and 14) for: GMEs (cyan dots), PREs (grey dots), and background regions of the genome (black dots). Horizontal lines show respective averages. **D. Genes associated with GMEs are enriched for functions in embryonic neuronal development:** Proportions of GME-associated genes (864 genes annotated, 90% with functional information) that fall into the subsequent categories: neuronal genes (72% versus 53.6% whole genome), neuro-developmental genes (60% GME versus 32.2% whole genome) and embryonic neuro-developmental genes (53% GME versus 27.2% whole genome). Genes and flybase keywords are listed in **supplemental data 1**. **E. GME-GME Interactions in Neuroblasts:** *Left*, Schematic of an Hi-C matrix and average Hi- C submatrices at GME-GME interaction. The Hi-C matrix contact intensity is visualized as a gradient from yellow (high) to green (medium) to blue (low). TADs are defined as domains of enriched contact, in various levels of green. GME positions are shown as vertical lines (cyan). Strong contacts are shown in yellow. For average Hi-C submatrices, all-to-all GME interaction submatrices were averaged. *Right*, Average Hi-C submatrices of all possible GME-GME interactions in cis (intra chromosomal arms, from 0Mb to 32Mb) and trans (inter arms and chromosomes) from KR-corrected and Observed/Expected transformed 25 kb resolution neuroblast Hi-C data (embryonic stage 10 and 14 combined). GME- containing Hi-C bins are pointed by black arrows. Total number of GME-GME interactions are shown in white.

### GMEs strongly associate with each other genome-wide and across long distances

We next performed *in situ* Hi-C to profile genome-wide chromatin contacts to examine whether GMEs are organized in the neuroblast genome. We purified neuroblasts using Fluorescence Activated Cell Sorting (FACS) from tightly-staged collections of a *dpn-eGFP* reporter embryo, which drives GFP from a minimal neuroblast enhancer of the *deadpan* (*dpn*) transcription factor gene (Awasaki et al., 2014; Lucas et al., 2021). Multiple batches of crosslinked neuroblasts were pooled for individual Hi-C experiments (**Supp Data 4**). Whole genome Hi-C matrices were generated using HiCExplorer at 25 kilobase (kb) resolution. Biological replicates were confirmed for reproducibility and quality control, following ENCODE criteria (Aiden, 2018), and we found that 75% of the TAD borders showed strict overlap, indicating high reproducibility (Sauerwald et al., 2020) (**Supp Fig. 2A**). Consistently, long-range contacts between Hox clusters that are 10Mb apart were similarly identified in individual replicates (**Supp Fig. 2B**). To examine GME-based interactions across the genome, we generated a megamap of aggregate reads from all replicates, producing an Hi-C matrix of 97 million contacts. We generated submatrices averaging Hi-C contact data from all possible pair-wise interactions between GMEs (**Fig. 1E**, schematic). GMEs are smaller than the Hi-C resolution, ranging from 68bp to 2.2 kb (average: 458bp ± 286bp). We thus refer to interactions between 25 kb bins containing GMEs as “GME interactions.” We profiled GME interactions by looking at the Observed versus Expected (Obs/Exp) values, which reveals distance-independent genome interactions. We found that GMEs interact across the genome, both in *cis* and in *trans* (**Fig. 1E**), and the specificity and frequency of interactions are robustly detected, even upon removing effects from strong contacts formed from short genomic distances (**Supp Fig. 2D**). We measured the average submatrices of background regions at 100 kb, 500 kb, and 1Mb distances from the GME contact site, further supporting the specificity of the GME interactions (**Supp Fig. 2C**).

**Figure 2:**
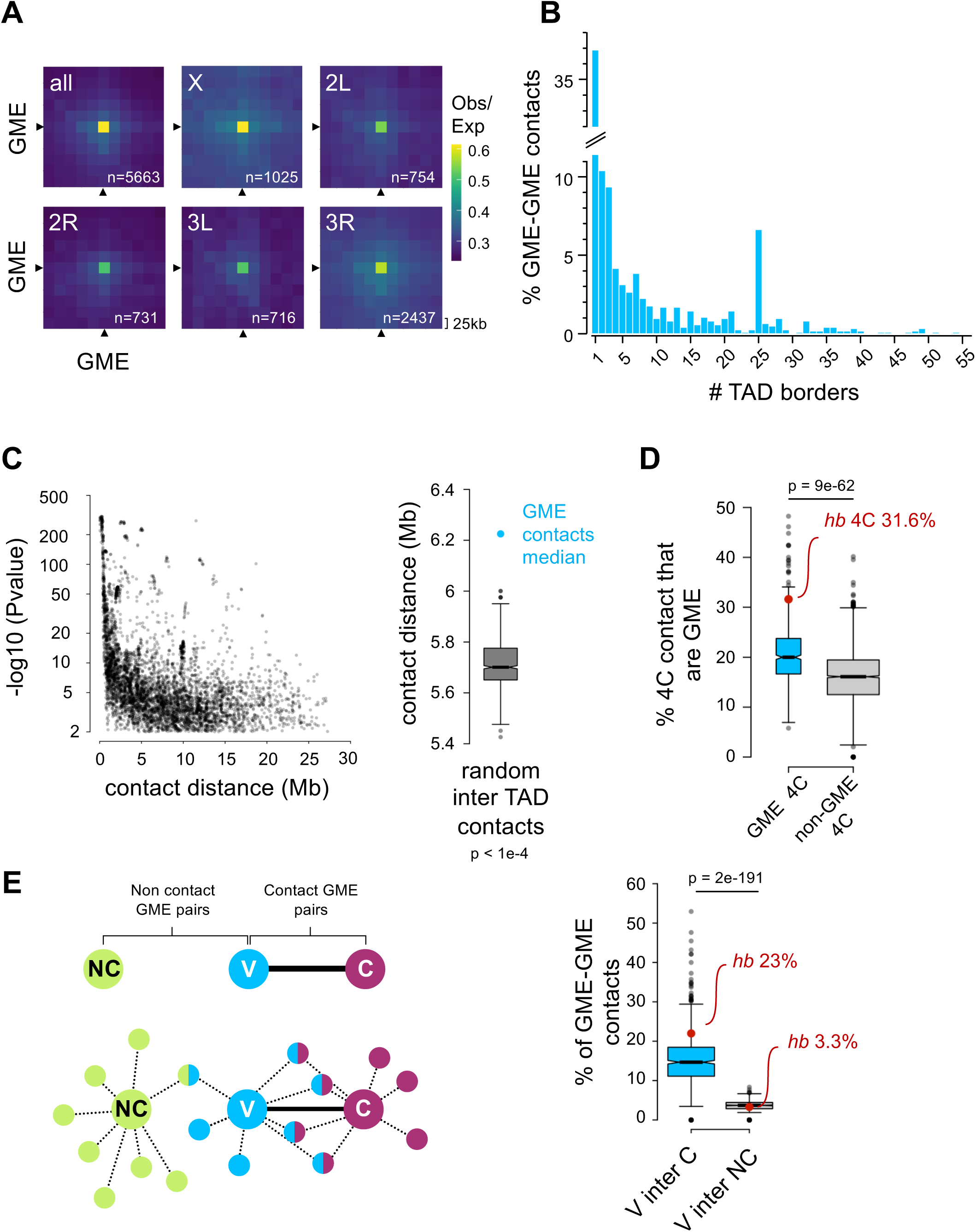
In neuroblasts, GMEs makes significant long-range contacts. A. GMEs make contacts genome-wide in neuroblasts: Average Hi-C submatrices of cis GME- GME significant contacts (p < 0.01) are shown by chromosome arms. GME- containing Hi-C bins are pointed by black arrows and the total number of GME-GME contacts (n=) is shown in white. 25 kb resolution Hi-C matrices were KR-correct and the Observed/Expected (O/E) is shown. **B. GME-GME contacts span multiple TAD borders:** Proportion of GME-GME significant contacts in neuroblast Hi-C data, distributed as a function of the number of TAD borders crossed. **C. GMEs make particularly long range contacts:** *Left*, Distribution of p-values as a function of contact distance for GME-GME contacts across TADs. Right: Boxplot comparing the median contact distance of GME-GME inter-TAD contacts (cyan dot) to the distribution of median contact distance from randomly sampled significant inter-TAD contacts. 10000 random sampling were performed (bootstrap p-value < 1e-4). The black bar represents the median of the random distribution. **D. Virtual 4C shows GMEs contact other GMEs more than non-GMEs:** This analysis was performed by extracting the 4C network of individual viewpoints from the Hi-C data. Quantification of the proportion of GME found in the virtual 4C of individual viewpoints, depending whether the viewpoint is a GME itself (“GME viewpoint”, cyan) or not (“non-GME viewpoints”, grey). The median proportion of GME in GME 4C and non-GME 4C is 20% and 16% respectively. Wilcoxon test was used for significance. **E. GME clustering within chromatin networks:** *Left*, Schematic illustrating GME clustering in 4C networks. The 4C network of all individual GME viewpoints is generated and the proportion of contact overlap between every 4C network is calculated. In this schematics, the 4C network of three GMEs are shown: in cyan, V (the <u>v</u>iewpoint GME); in purple, C (a GME that is in <u>c</u>ontact with V); and in green, NC (a GME that is <u>n</u>ot in <u>c</u>ontact with V). Each of these GMEs in this example all make a total of 8 contacts with other GMEs. GMEs that belong to two networks are bi-color. Note that there are more shared contacts between V and C than between V and NC (4/8 vs. 1/8 respectively). *Right:* Quantification of the proportion of contacts shared between the 4C networks of two GMEs that are in contact (V inter C) or between the 4C networks of two GMEs that are not in contact (V inter NC) . Median proportion of shared GME contacts in “V inter C” and “V inter NC” are 14.6% and 3.7% respectively. For *hb* GME viewpoint the proportion of shared GME contact in “V inter C” and “V inter NC” is 23% and 3.3% respectively. Wilcoxon test was used for significance.

We next implemented FitHiC2 (Kaul et al., 2020) and HiC-ACT (Lagler et al., 2021) to detect statistically significant chromatin interactions (contacts) at 25 kb resolution. We found that distribution of the contacts as a function of Hi-C read counts had a median of ten contacts and were largely similar between GME contacts and all chromatin contacts (**Supp Fig. 3A**). Further, GME interactions were strongly detected in individual chromosome arms, indicating that GME interactions exist genome-wide (**Fig. 2A**). Finally, we found that GMEs formed contacts that crossed TAD boundaries (median 6.2Mb), with nearly 40% of GME interactions crossing at least one TAD border, and many crossing multiple TAD boundaries (**Fig. 2B, Supp Fig 3B,C**). These contacts were highly statistically significant (**Fig. 2C**) and included GME interactions even over 20Mb distances. Notably, we found that inter-TAD GME interactions were detected over significantly longer distances compared to regions that do not contain GMEs (**Fig. 2C**). In fact, 94% of all *cis* GME interactions were inter-TAD contacts (**Supp. Fig 3B**), consistent with widespread GME distribution across the genome, but with very few GMEs within individual TADs. These data indicate robust, long-range GME contacts in neuroblasts.

**Figure 3:**
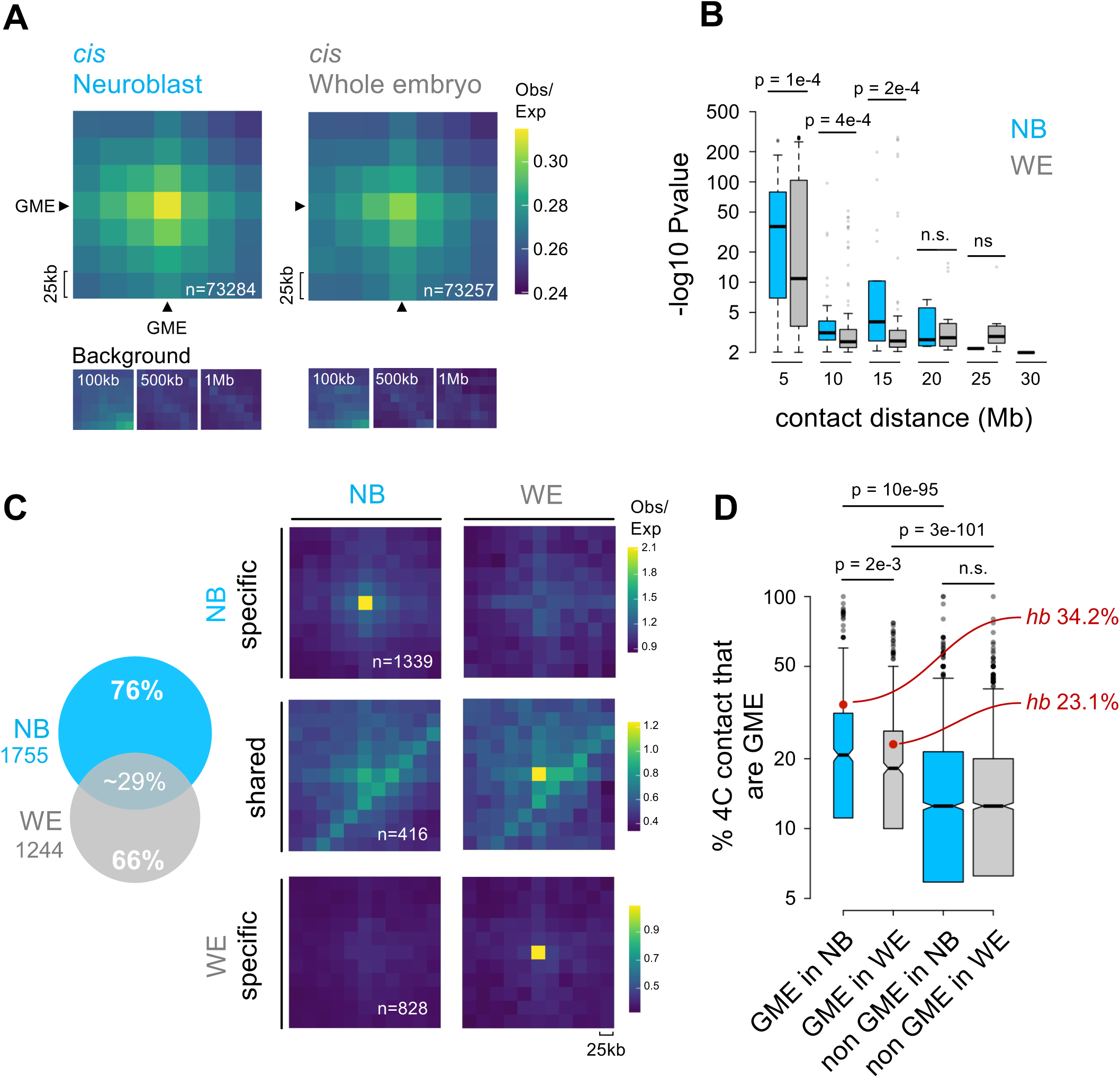
GMEs form cell type specific interactions. A. GME interactions are stronger in neuroblasts than in whole embryo: *top*: Average Hi-C submatrices of all possible GME-GME cis interactions in neuroblast and whole embryo cells. 25 kb resolution HiC matrices were normalized, KR-corrected and Observed/Expected values are displayed. GME-containing Hi-C bins are pointed by black arrows. Total number of GME-GME interactions are shown in white. *Bottom:* Average submatrices, sampled at distances of 100 kb, 500 kb, and 1 Mb from the GME-GME interaction. **B. GMEs make more long-range contacts in neuroblasts than in whole embryo:** Distribution of p-value as a function of contact distance (5Mb bins) in neuroblast (cyan) and whole embryo (grey). Wilcoxon test was used for significance. **C. Most GME-GME contacts are cell type specifics:** *Left*, Venn diagram of all cis and trans GME-GME significant contacts detected in neuroblast (cyan) and whole embryo (grey) *Right:* Average Hi-C submatrices of neuroblast specific, whole embryo specific and shared significant GME-GME contacts in neuroblast and whole embryo. Hi-C data were normalized, KR-corrected and Observed/Expected values were displayed. All contact distances are shown (From 0Mb to 32Mb). Number of contacts (n=) are shown in white. **D. 4C analysis of GME contact propensity:** Quantification of the proportion of cis and trans GME-GME contacts in virtual 4C for each GME viewpoint and non-GME viewpoints in neuroblast (cyan) and whole embryo (grey). Median shown as black horizontal line. Wilcoxon test was used for significance.

We next used the Hi-C matrix to generate virtual 4C interactomes (4C) and compared the 4C of GME-containing bins with the 4C of bins that do not contain GMEs. We found that compared to non-GMEs, GMEs had greater affinity for other GMEs (**Fig. 2D**). GMEs made on average 192 contacts, of which 20% were with other GMEs (**Supp Fig. 4A**). The *hb* GME was enriched for GME contacts (*cis* and *trans*) at 31.6% (**Fig 2D**, red dot, and **Supp Fig. 4A**), and this was a significant enrichment above the 12% of GME-containing bins genome-wide (Hi-C bins containing GMEs = 665; total Hi-C bins = 5505). Further, the *hb* GME interaction with other GMEs was stronger than with non-GME sites (**Supp Fig 4B, inset**). The specificity of the interactions is exemplified by the *hb* GME interaction with the *BX-C* locus, which shows at 25 kb bin stripe across the 300 kb locus (**Supp Fig. 4B**).

**Figure 4:**
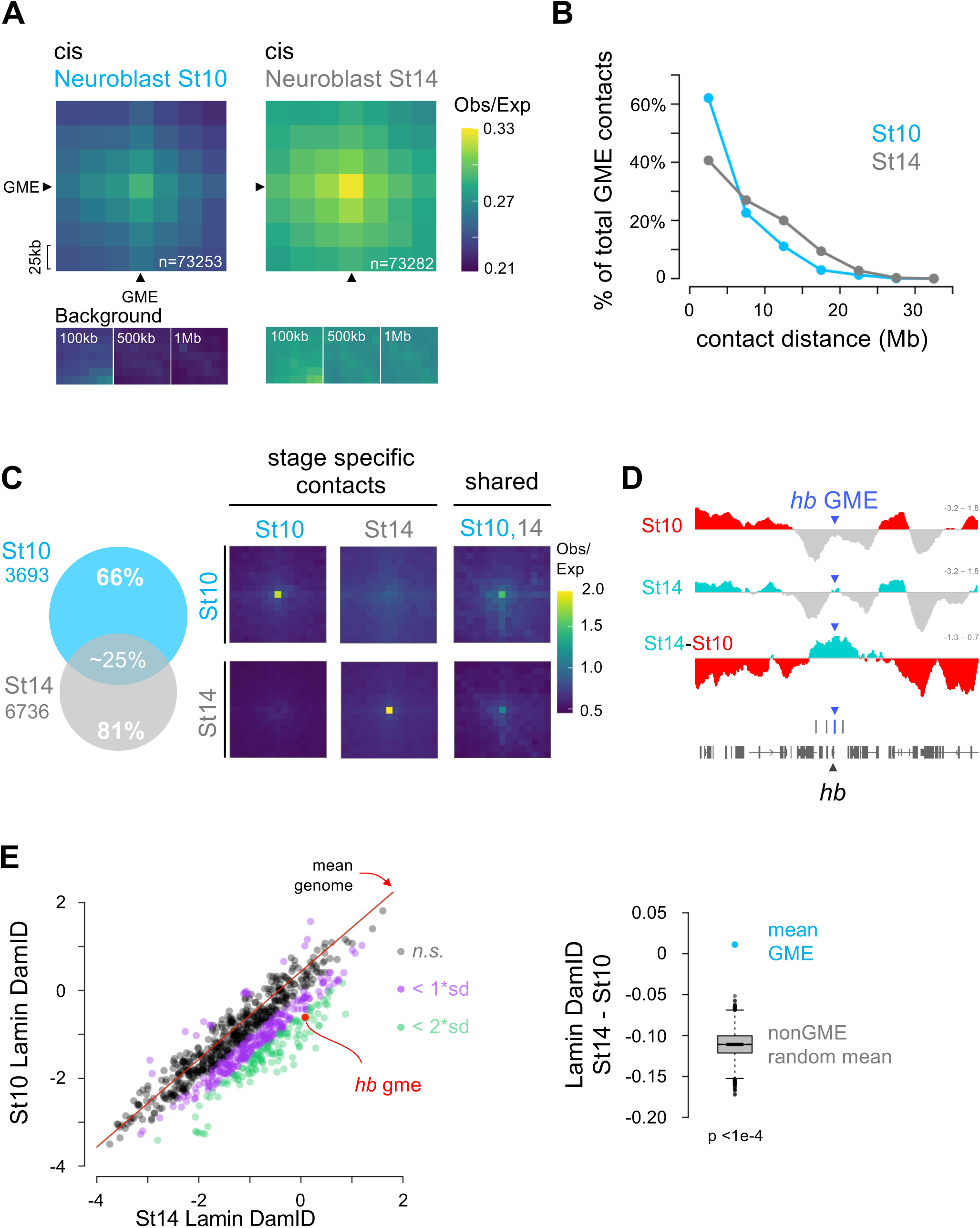
GMEs form temporal specific interaction in neuroblast. A. Cis GME-GME interactions are stronger over developmental time: *Top*, Average Hi-C submatrices of all possible GME-GME cis interactions in embryonic stage 10 and stage 14 neuroblasts. 25 kb resolution HiC matrices were stage normalized, KR-corrected and the Observed/Expected values are displayed. GME-containing Hi-C bins are pointed by black arrows and the central bin represents the GME-GME interaction. Total number of GME-GME interactions are shown in white. *Bottom:* Average submatrices, sampled at distances of 100 kb, 500 kb, and 1 Mb from the GME-GME interaction. **B. GMEs make longer-range contact in neuroblasts stage 14 compared to stage 10:** The proportion of significant cis GME-GME contacts is plotted as a function of contact distance (5 Mb bins). Stage 10 and stage 14 are shown in cyan and grey, respectively. **C. Most GME-GME contacts are stage specifics:** *Left*, Venn diagram of cis and trans GME- GME significant contacts detected in neuroblast stage 10 (cyan) and stage 14 (grey). *Right,* Average Hi-C submatrices of neuroblasts stage specific and shared GME-GME contacts. Hi-C data were normalized, KR-corrected and Observed/Expected values were displayed. All contact distances are shown (0 Mb to 32 Mb). **D. The frequency of *hb* GME interaction to the lamina is higher in neuroblasts stage 14 compared to stage 10:** Lamin DamID profiles spanning the *hunchback* gene region at both stages 10 and 14 are presented as log₂ ratios of Dam::Lamin signal over Dam signal. Positive signals are shown in red and turquoise for stage 10 and stage 14, respectively. Differential Lamin DamID profile is shown as the difference between the stage 14 and stage 10 Lamina DamID profiles, with red indicating higher frequency at the lamina at stage 10 and turquoise indicating increased frequency at the lamina at stage 14. The positions of GMEs and genes are marked as blue bars and gray blocks, respectively. **E. The frequency of GMEs interaction with the lamina is higher in stage 14 neuroblast compared to stage 10 and compared to the average frequency of interaction to the lamina for non-GME loci::** *Left,* Scatter plot showing the stages 10 and stage 14 Lamin DamID signals for each GME. The red diagonal represents the average of the differential Lamin DamID (stage 14 - stage 10) genome-wide. GMEs with stage-specific differences exceeding 1 or 2 standard deviations above the genome-wide average are highlighted in purple and green, respectively. The stage-specific DamID signal at the *hunchback* GME is indicated in red. *Right,* Boxplot comparing the average of the differential Lamin DamID of GME (cyan dot) to the distribution of average of differential Lamin DamID from the random sampling of nonGME ATACseq regions genome-wide. 10000 random sampling of 859 ATAC sites were performed (bootstrap p-value < 1e-4). The black bar represents the median of the random distribution.

Finally, we found that GMEs appear to show some level of clustering (**Fig. 2E**). We generated the virtual 4C of each GME and calculated the proportion of contact overlap between every 4C network. For all possible pairwise 4C viewpoints, we separated pairs that interact (<u>V</u>iewpoint GME interaction with <u>C</u>ontacting GME, “V-C”) from pairs that do not interact (<u>V</u>iewpoint GME interaction with <u>N</u>on-<u>C</u>ontacting GME, “V-NC”). We found substantially greater shared GMEs among the 4C network between two GMEs that directly interact (**Fig. 2E**, blue-pink in schematic) than between two GMEs that do not interact directly (**Fig. 2E**, green-blue in schematic). Taken together, we conclude that GMEs preferentially interact with other GMEs, and there appears to be clustering among GME subsets.

### GMEs interactions are cell type specific

A central question in the field of genome architecture is how genome organization reflects cell type specificity (Alexander and Lomvardas, 2014; Bashkirova et al., 2023; Lucas and Kohwi, 2019; Tan et al., 2019; Winick-Ng et al., 2021). To test whether there are aspects of GME-based genome organization specific to neuroblasts, we generated Hi-C datasets from whole embryos to compare to our neuroblast Hi-C datasets generated above. To this end, we FACS-purified live cells from stage-matched, whole embryos using the same *dpn-eGFP* reporter genotype from which we sorted neuroblasts, and we obtained comparable numbers of cells as that which were fixed and processed for neuroblast low-input Hi-C (**Supp data 4**). A recent study on whole embryo single cell RNA sequencing suggests our live cell mixture contains epidermal, muscle, and other cell types, with nervous system cells constituting 10-15% (Peng et al., 2024). Correlation analyses of the Hi-C dataset show that while the level of correlation is high overall (>0.8), neuroblast (NB) replicates correlated more with each other than with whole embryo (WE) replicates (**Supp Fig. 5A**). NB and WE replicates Hi-C data were pooled, normalized together and KR corrected, generating 25 million contact maps. We profiled and compared Obs/Exp values of GME interactions in both cell types, and we found that *cis* GME interactions were stronger in NB than in WE. (**Fig. 3A, Supp Fig. 5B).** Using FitHiC2 followed by HiC-ACT pipelines on the normalized NB and WE Hi-C data to detect significant GME contacts, we found that when GME contacts were binned by contact distance (i.e. genomic distance between GMEs), NB generated more significant GME contacts than in WE across long distances (**Fig. 3B**), and the GME contact *P* values were also generally stronger **(Supp Fig. 5C**).

**Figure 5:**
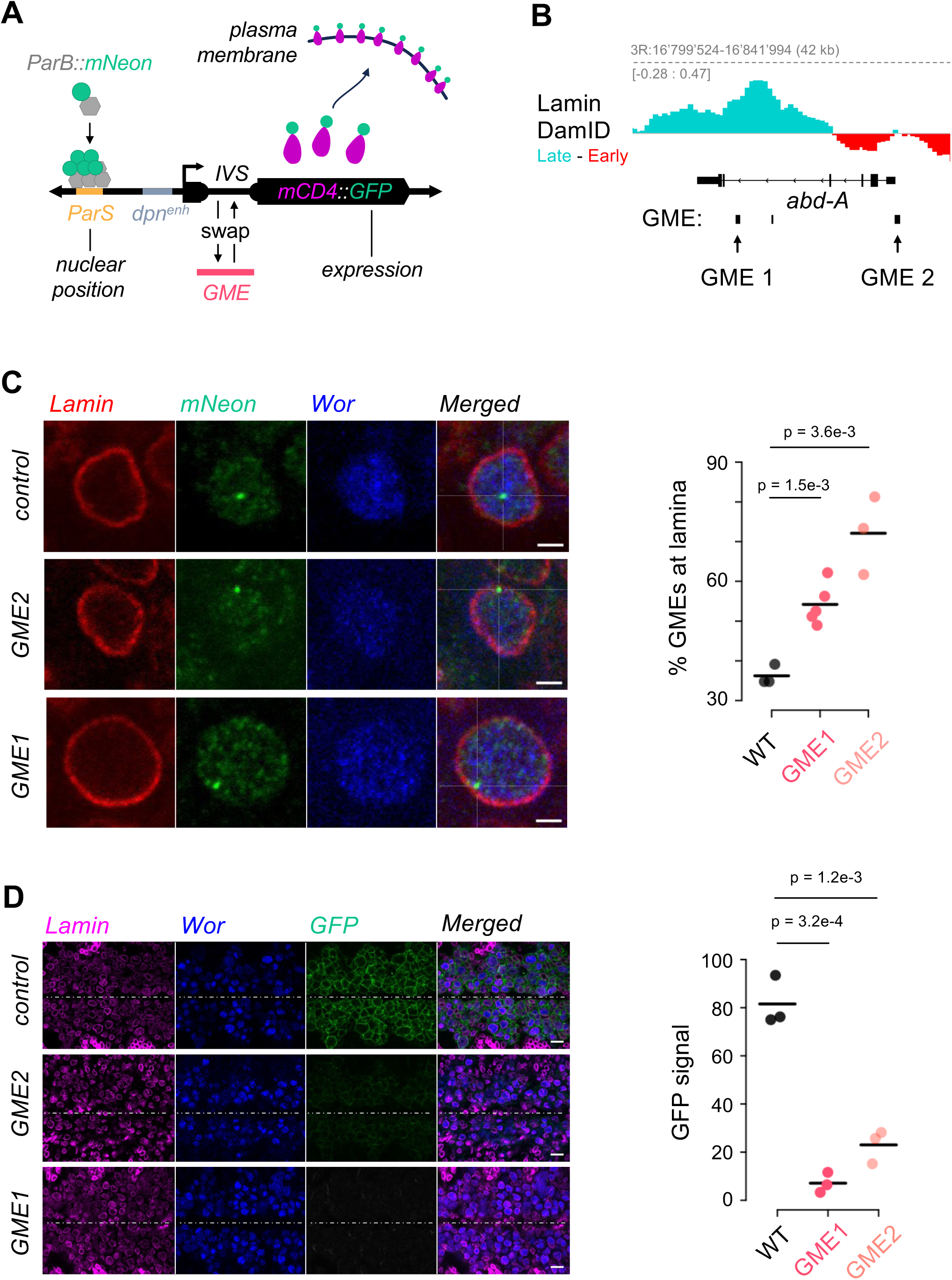
*in-vivo* validation of GME function using *abd-A* GMEs **A. Design of the GME reporter transgene:** Schematic displaying the structure of the ParS- dpn>{IVS}-mCD4::GFP reporter transgene. The construct combines nuclear positioning (ParS/ParB::Neon) and gene expression markers (mCD4::GFP). mCD4::GFP expression is driven by the *dpn* neuroblast enhancer (dpn^enh^), with the IVS synthetic intron replaced by *abd-A* GME1 or *abd-A* GME2 sequences. **B. Both *abd-A* GMEs interact with the lamina more frequently in stage 14 neuroblasts than in stage 10:** stage 14 - stage 10 differential Lamin DamID tracks in neuroblasts at the abd- A locus. The signal is colored in red when the frequency of interaction to the lamina is greater at stage 10 and it is colored in turquoise when it is greater at stage 14. **C. Both *abd-A* GMEs individually promote nuclear lamina localization of the reporter:** *Left:* Representative neuroblast single z-plane images at stage 14 showing Lamin (nuclear envelope), ParB-Neon (transgene locus), and Worniu (neuroblast marker) for IVS control and GME reporters. Scale bars, 2 μm. *Right:* Quantification of loci localized at the nuclear lamina, showing increased lamina association for GME1 and GME2 constructs compared to the control (IVS) in stage 14 neuroblasts. Statistics: t-test. **D. Both *abd-A* GMEs individually repress mCD4::GFP expression in neuroblasts:** *Left:* Single z-plane images of the ventral nerve cord neuroblast layer at stage 14 for IVS (WT), GME1, and GME2 constructs, stained for Lamin (magenta), Worniu (blue), and mCD4-GFP (green). Midline indicated by dashed line. Scale bars, 10 μm. *Right:* Quantification of normalized GFP fluorescence intensity across three embryos per condition, showing GME-mediated repression of reporter expression. Statistics: t-test.

To determine the level of cell type specificity of GME contacts, we next compared the significant GME interactions in NB and WE, and we found that while there was significant overlap of TAD borders between the two (**Supp Fig. 5D**), 76% of the GME contacts found in NB were not significant in WE sample (**Fig. 3C** and **Supp Fig. 5E**). In fact, when we analyzed the virtual 4C for each GME (one GME viewpoint to all contacts), we found that compared to GMEs in WE, GMEs in NB were more enriched for GME contacts. This was in contrast to Hi-C bins that do not contain GMEs, in which the fraction of GME contacts they made were comparable between NB and WE. (**Fig. 3D**). Notably, we analyzed Hi-C datasets from a recent study on *Drosophila* chromatin organization in the nervous system (Mohana et al., 2023), and found that GME interactions were stronger in neuroblasts than in glial cells (**Supp Fig. 5F**). Taken together, the results show that GMEs reveal a cell type specific genome organization marked by a robust and enriched interaction in neuroblasts.

### GME organization is dynamic over time and mediates gene mobility to neuroblast nuclear lamina in late stage embryos

Neuroblasts produce different neural cell types over time, and we previously showed that a mid- embryonic relocation of the *hb* gene to the nuclear lamina, mediated by its intronic GME, terminates competence to produce early-born neurons (Lucas et al., 2021). Thus, we explored whether the GME interactions are dynamic. To this end, we compared GME contacts in Hi-C normalized data from stage-specific neuroblasts, stage 10 and 14 (before and after the stage when the *hb* gene relocates to the nuclear lamina). We found that while the TADs are largely the same, with 72% of the TAD borders overlapping between the early and late stages (**Supp. Fig. 6A**), GME interactions changed. The Obs/Exp averages of pairwise *cis* GME interactions were higher in older neuroblasts compared to younger neuroblasts (**Fig. 4A, Supp Fig. 6B,C**). Further, we found that the distribution of significant (*P* value<0.01) GME contacts across distances were consistently higher in older neuroblasts compared to younger (**Fig. 4B**). Notably, when we analyzed the stage-specific GME contacts, we found that 66% of the GME contacts detected at stage 10 and 81% of GME contacts detected at stage 14 were stage-specific interactions (**Fig. 4C, Supp Fig. 6D**). The stage 14 specific GME contacts were longer-range and had greater statistical significance than the stage 10 specific contacts (**Supp Fig. 6E**). Together, these data indicate that while GMEs maintain enriched interactions with other GMEs, the specific GME partnerships are dynamic over time.

Finally, in addition to changes in chromatin contacts, we examined whether genomic sites containing GMEs relocate to the neuroblast nuclear lamina like the *hb* gene. We performed *in vivo* targeted Lamin:DamID, a DNA-methylation-based technique to map genomic regions localized at the nuclear periphery (Aughey and Southall, 2016; Southall et al., 2013; van Steensel and Henikoff, 2000). To transiently drive Dam-fused Lamin expression specifically in neuroblasts, before and after the mid-embryonic genome reorganization, we used the UAS/GAL4 system, via heat-shock using the thermosensitive GAL4 inhibitor GAL80^ts^ (Southall et al., 2013). We compared lamina-associated genes in neuroblasts at stage 10 or stage 14, and first confirmed that the *hb* GME increases its contact frequency with the nuclear lamina at stage 14, consistent with our *in vivo* observations (Kohwi et al., 2013; Lucas et al., 2021) (**Fig. 4D**). Next, by taking the difference in lamina association between stage 14 and stage 10 (Differential DamID), we identified regions that increase or decrease their frequency of lamina association genome-wide, indicative of gene mobilization. We found that relative to the average differential DamID value of the whole genome (**Fig. 4E**, red diagonal line, **Supp Fig. 7A**), GMEs as a group significantly increased Lamin Dam signals in late stage neuroblasts, indicating a greater association with the nuclear lamina (**Fig. 4E right and Supp Fig. 7B**). Further, we found that the differential DamID signal of GMEs was significantly greater than PREs (**Supp Fig. 7C**) or the recently described tethering elements (TEs) (**Supp Fig. 7D**), shown to facilitate enhancer-promoter interactions in the early embryo by mediating distal chromatin interactions (Batut et al., 2022; Levo et al., 2022). Consistently, we did not observe substantial overlap of GMEs with either PREs or TEs. Thus, despite their common characteristic of being associated with PcG chromatin factors, they appear to have distinct features in context of genome organization in the neuroblast (see Discussion).

### GMEs mediate gene relocation in neuroblasts

To test the ability of newly discovered GME sequences to relocate to the lamina *in vivo*, we generated transgenic flies in which we swapped the synthetic intron of a GFP reporter transgene for a GME sequence. The mCD4::tdGFP reporter drives expression of a plasma-membrane- localized GFP under the control of the neuroblast-specific *deadpan* promoter (Awasaki et al., 2014). The construct also includes a *ParS* bacterial sequence that allows us to observe the transgene’s physical position in the neuroblast nuclear space through detecting a ParB::mNeon fusion protein that is recruited of the to the *ParS* sequence (Saad et al., 2014) (**Fig. 5A**). As proof of principle, we selected two candidate GMEs that share similar characteristics to the *hb* GME, including a positive change in Lamin:DamID signal between early and late stage embryos. GME1 is in the intron of the *Abd-A* gene, and GME2 is near the promoter (**Fig. 5B**). Embryos were fixed and immunostained for mNeon, lamin Dm0, which marks the nuclear envelope, and Worniu, a pan-neuroblast marker. In control animals, the unmodified reporter transgene was localized in the nuclear interior, whereas in experimental animals, in which the intron was replaced with either GME1 or GME2, showed a significant increase in localization at the nuclear lamina (**Fig. 5C**). Given that the nuclear lamina is highly associated with gene silencing (Shevelyov and Nurminsky, 2012; van Steensel and Belmont, 2017), we measured the GFP expression levels of the reporter transgene with and without the GME sequence replacement. We quantitatively measured the mCD4-GFP signal by sampling 100 random positions at the neuroblast plasma membrane and subtracted background signal taken from 50 random background regions (outside of the cells) (**Supp Fig. 8A)**. We found that while the control GFP reporter is transcriptionally active (GFP^+^), both GME1 and GME2-swapped transgenes showed markedly lower GFP expression levels, consistent with their transcriptional silencing at the nuclear lamina (**Fig 5D** and **Supp Fig. 8B)**;These data are highly reminiscent of the *hb* GME, which also resulted in lamina localization of a GFP reporter transgene and a concomitant reduction in GFP expression (Lucas et al., 2021). Together, the data show that GMEs mediate lamina-association *in vivo*.

## DISCUSSION

While great progress has been made in revealing the fundamental principles of genome architecture at multiple scales, what level of organization relates to gene function is still largely unknown. Challenges include limited data that take into account the high context-specificity of genome architecture, which require *in vivo*, cell type specific, and developmental stage-specific analyses. Much attention on genome architecture has focused on TADs, but how TADs relate to functional genome organization remains controversial. Genomic interactions within TADs were found to be only marginally stronger than between TADs (Finn and Misteli, 2022), and microscopy studies show that TADs are emergent statistical properties among populations of cells each with dynamic chromatin conformations (Bintu et al., 2018; Chi et al., 2022). Biochemical mechanisms have been proposed to better link TAD-based chromatin interactions with gene expression (Xiao et al., 2021). Nonetheless, global loss of TAD boundaries leads to minimal dysregulation of overall gene expression (Ghavi-Helm et al., 2019; Nora et al., 2017; Rao et al., 2017; Stik et al., 2020), and widespread changes in transcription does not necessarily correlate to major changes in genome-wide chromatin organization (Ing-Simmons et al., 2021). Thus, what kind of genome organization impacts the cell’s functional output is still an open question.

Here we chose to approach genome organization from a reductionist perspective, by starting with a regulatory element known to mediate biologically relevant gene reorganization (Kohwi et al., 2013; Lucas et al., 2021). We explored whether similar sites exist genome-wide and whether they are subject to cell type and stage specific organization. We previously showed that the *hb* gene undergoes a developmentally-timed relocation to the nuclear lamina in the neuroblast progenitor that closes the competence period to produce descendent neurons that can transcribe the *hb* gene endogenously (Kohwi et al., 2013), a molecular marker of early-born identity. We identified an intronic *cis*-regulatory element that is both necessary and sufficient for its mobility to the nuclear lamina. One of the key features was that this element (GME) maintained chromatin accessibility even after its relocating to the nuclear lamina and the rest of the gene body and the promoter became chromatin inaccessible, consistent with its heritably silenced state. Further, loss of PRC1 reduced *hb* gene localization at the neuroblast nuclear lamina without concomitant transcriptional derepression (Lucas et al., 2021). Thus, we sought to identify other genomic sites, prioritizing high chromatin accessibility and that are PcG target sites. An elegant study relating PcG activity to nuclear pore complex proteins suggest that a subset of PcG target genes are silenced at the nuclear envelope through coordinated activity of Nup93, indicating that PcG target genes may be selectively and differentially regulated through co-partners of PcG factors (Pascual- Garcia et al., 2017). Thus, using these criteria, we identified over 800 putative GME sites across the genome first, and then asked whether and what kind of organization is revealed. We found that GMEs revealed a robust network of interactions in neuroblasts that would not have been apparent from a genome-wide Hi-C map. We found that GME interactions make long-distance interactions that cross multiple TAD boundaries, and while GMEs preferentially interact with other GMEs, their specific GME partners were both cell type and stage-specific, suggesting a high context specificity to the GME interacting network.

Notably, we found that as a group, GMEs mobilize to the neuroblast nuclear lamina in late stage embryos, and swapping a control transgene intronic sequence with a GME was sufficient to increase localization of the transgene at the neuroblast nuclear lamina and reduce GFP expression. The nuclear lamina is established as a subnuclear domain highly associated with silenced or repressed genes (Shevelyov and Nurminsky, 2012; van Steensel and Belmont, 2017). Our previous work with the *hb* gene showed that an already transcriptionally repressed gene can become further heritably silenced upon relocation to the nuclear lamina. In fact, CRISPR- mediated deletion of the *hb* GME did not result in any changes in *hb* transcriptional dynamics – *hb* turned on and off on time. Rather, the GME deletion impacted only *hb* gene relocation to the neuroblast nuclear lamina, and consequently, its heritably silenced state. Thus, while the nuclear lamina is typically associated with transcriptional silencing, it can also act in epigenetic regulation, and the consequences on the gene upon relocation is context-dependent (Lucas et al., 2021). Indeed, when we swapped the GFP reporter transgene intron with a GME, the transgene increase association with the nuclear lamina and GFP expression was reduced. Perhaps GMEs function to facilitate the process of genome reorganization while remaining agnostic to the regulation of individual gene transcription. Other studies have identified regulatory elements that function in genome organization in context of transcriptional regulation. Subsets of PREs have been found to mediate long-range chromatin contacting to facilitate PcG-mediated silencing (Bantignies et al., 2011; Ogiyama et al., 2018), and PRC1 proteins have been shown to play a role in chromatin compaction (Cheutin and Cavalli, 2018). Tethering elements (TEs) have been identified as regulatory sites that facilitate communication between distal enhancers and promoters to mediate transcription, without necessarily functioning as enhancers themselves (Batut et al., 2022; Levo et al., 2022). TEs are also bound by PRC1 proteins, and a subset of TEs have also been identified as PREs (Denaud et al., 2024; Li and Levine, 2024). Both PREs and TEs are PcG targeted genomic sites, which is also true for the GMEs we identified in this study. There appears to be some, but not substantial overlap between GMEs and PREs or TEs. Notably, PREs, based on a machine-learning algorithm that predicted the majority of PREs from multiple experimental datasets (Bredesen and Rehmsmeier, 2019), were distinct from GMEs in that they were largely chromatin inaccessible in neuroblasts. When we compared the stage-dependent mobility of these elements relative to the neuroblast nuclear lamina in our Lamin:DamID dataset, we found that GMEs significantly increased association with the lamina compared to either PREs or TEs. All of these elements have PcG targeting in common, and yet their accessibility and behaviors differ. On a complementary note, PcG chromatin factors are ubiquitously expressed, and yet, their regulatory functions are highly cell and stage specific. It is exciting to speculate that PcG chromatin factors may have multiple, functionally distinct, types of target sites on the genome that are differentially recruited, perhaps through cell type specific chromatin accessibility, to confer unique functions in the cell. We also note that neuroblasts are markedly devoid of the H3K27me3 histone mark associated with PRC2-mediated repression, similar to the early embryo stage when TEs were found to mediate chromatin contacts (Batut et al., 2022). Moreover, a study of the *vestigial* gene revealed two PREs, one near the promoter and another intronic one, with distinct functions on the gene (Ahmad and Spens, 2019). In our GME dataset, only the promoter PRE, but not the distal PRE that contributes to H3K27me3, is a GME and is characterized with Nup93 ChIPseq peak. Perhaps these are examples in which PcG function is utilized for genome organizational functions that may be distinct from transcriptional repressive activities through H3K27me3. These are indeed fascinating questions to be addressed in the future.

## Data availability

The datasets generated during and or/analyzed during the current study are available from the corresponding author upon reasonable request.

## Acknowledgements

We thank the members of the Kohwi lab for their scientific discussion and feedback. We thank Raphaelle Laureau for feedback on the manuscript. This work was funded by the Hirschl Trust Foundation, NIH R01HD092381 and R01GM152663 to M.K.

## Author Contributions

T.L. and M.K. designed the experiments, analyzed the data, and wrote the manuscript. T.L. performed the Hi-C experiments, immunostaining, and bioinformatic analyses. L-I.W. performed the Lamin DamID experiments. N.M. performed the FACS and generation of sequencing libraries. E.Q. determined the genes associated with GMEs. J.G.K. performed Hi-C analyses and contributed to manuscript writing. S.P. generated the GFP reporter constructs.

## MATERIALS AND METHODS

### Fly Lines

Wild-type *(w1118)*, *esc^6^* and *esc^5^* (Struhl, 1981; Tie et al., 1998), DamID related flies: *attp2{UAS- LT3-mCherry::Dam::myc::LaminDm0}*, *attp2{UAS-LT3-mCherry::Dam::myc}*, and *worGal4 ° tub- Gal80ts* (Southall et al., 2013); Dpn reporter flies: *dpn-DSCP-IVS-mCD4::GFP*, *dpn-DSCP- ParB::mNeon*, and *dpn-eGFP* (Lucas et al., 2021). Flies were raised on a standard cornmeal/molasses medium at 25C.

### Plasmids

The control transgene *ParS-dpn-DSCP-IVS-mCD4::tdGFP* (IVS plasmid) (Lucas et al., 2021) was modified to generated additional transgenes reported in this paper by replacing the *IVS* intron with *abd-A* GMEs sequences: *ParS-dpn-DSCP-{abd-A-GME1}-mCD4::tdGFP* and *ParS-dpn-DSCP-{abd-A-GME2}-mCD::tdGFP*.

### Antibodies

Rat monoclonal Anti-Worniu (5A3AD2) (Abcam, ab196362) 1:100; Rat monoclonal Anti-Deadpan (11D1CH11) (Abcam, ab195172) 1:100; Chicken polyclonal Anti-GFP (Aves Labs, GFP1010, RRID: AB_2307313) 1:500; Rabbit polyclonal anti-Lamin R836 (Kind gift from Dr. Paul Fischer, Stony Brook University) 1:1000; Mouse anti-mNeonGreen, (32F6) (ChromoTek,32f6, RRID: AB_2827566) 1:500; Mouse monoclonal anti-Abd-B (1A2E9) (DSHB, 1A2E9, RRID: AB_528061) 1:100; Rabbit polyclonal anti-H3K27me3 (Sigma-Aldrich, 07-449, RRID: AB_310624) 1:200; DAPI (Thermo Fisher Scientific, D3571) 1:5000

### Immunohistochemistry

Embryos were immunostained according to standard protocols (Rothwell, 2000). Briefly, embryos were dechorionated with 50% bleach, extensively washed in water, and fixed by rocking for 22min at room temperature in a 1:1 mixture of 4% formaldehyde in PEM buffer (0.1M Pipes, 1mM MgS04, 2mM EGTA) and N-heptane. Embryos were then devitellinized by vigorous shaking in a 1:1 mixture of methanol:heptane and washed three times with PBS before staining. Embryos were incubated in primary antibodies in PBS-0.1% Tween 20 (PBT) overnight at 4°C, secondary antibodies at room temperature for 1.5hrs.

### Neuroblast purification

*dpn-eGFP* embryos were collected for 1-2hrs, dechorionated, and immersed in halocarbon oil 27 (Sigma-Aldrich, H8773) to identify stage. Embryos older than stage 4 were removed by hand to ensure developmental stage uniformity and then aged at 18°C for 18hrs to stage 14 or at 29°C for 3hrs to stage 10 before mechanical dissociation in PBS (tissue culture grade) with 2% fetal bovine serum. Cells were incubated with Calcein Violet 450 AM (Thermo Fisher Scientific, 65- 0854-39) and 7-AAD (Thermo Fisher Scientific, 00-6993-50) to label live and dead cells, respectively, and sorted on a Beckman Coulter MoFlo Astrios. FACS was performed as previously described (Lucas et al., 2021), in which cells were purified based on viability, size, granularity and GFP fluorescence intensity. FACS gates were adjusted to select for single, round, and GFP- positive neuroblasts while excluding neurons that inherited GFP protein and that had formed neuroblast-sized clusters. Before each neuroblast collection, 100 cells were sorted and examined under brightfield and fluorescence microscopes to confirm the above criteria. For Hi-C, live neuroblasts and live GFP-negative cells were collected into 100ml of cold PBS and fixed as described (Tan et al., 2021). Briefly, tubes with sorted cells were filled with ice-cold PBS up to 1ml, and 133.3ul of 16% Paraformaldehyde (methanol-free, Fisher, 28908) was added to each tube. Cells were fixed for 10 min at room temperature on a rotator. 100ul of cold PBS-1%BSA (Jackson Immuno, 001-000-161) was added to each tube to quench PFA. After centrifugation for 5 min at 600xg (4°C), cells were washed once with PBS-1%BSA. Supernatant was removed, leaving 30ml in the tubes and stored at -80°C.

### ATAC-seq analysis

Neuroblast ATAC-seq for embryonic stages 10, 12 and 14 were generated in (Lucas et al., 2021). Data for stage 14 is newly published here and analyzed as described in (Lucas et al., 2021). Briefly, raw sequencing reads were converted to FASTQ using bcl2fastq (Illumina) and quality controlled using FASTQC (Wingett and Andrews, 2018). After adaptor trimming using cutadapt (- m5 -e 0.2 -o {R1} -p {R2}) (Martin, 2011), paired-end reads were aligned on dm6 using bowtie2 (- k 4 -X2000 –local –mm) (Langmead and Salzberg, 2012). Aligned reads were deduplicated and filtered for low quality reads using Picard (Picard Toolkit, BroadInstitute) and Samtools (Li et al., 2009), respectively. Read positions were shifted (+5/-4 bp) (Buenrostro et al., 2013). Finally, samples were “read per million” (RPM) normalized using BEDTools genomecov (Quinlan and Hall, 2010). ATAC-seq normalized tracks were visualized as bigwig file (Kent et al., 2010) on IGV (Thorvaldsdottir et al., 2013). Peaks were called using MACS2 (Gaspar, 2018) on ATACseq fragments smaller than 100bp that are representing open regions as described in Lucas et al., 2021.

### GME associated gene characterization

<u>Putative Gene Mobility Elements were extracted from the intersection of multiple peak lists:</u> ATACseq, Polycomb ChIPseq (GSE102339) (Brown et al., 2018) and Nup93 ChIPseq (GSE94922) (Pascual-Garcia et al., 2017) using intervene venn script (Khan and Mathelier, 2017). Peaks from neuroblast ATAC-seq embryonic stages 10, 12 and 14 were intersected and peaks present in at least two stages were kept. These ATAC peaks were subsequently intersected with the peaks detected in Polycomb (E(z), Psc and Pho) and Nup93 ChIP-seq datasets, resulting in the list of 859 putative GME regions. Peak detections for ATAC and ChIP datasets were performed using MACS2. MACS2 peak significance threshold (*P* values) for ATAC (St10 to 14), Nup93, Pho, E(z) and Psc were less than 1e-6, 1e-2, 5e-4, 5e-4 and 5e-2, respectively. Each of the peaks in the ChIP-seq peak list were visually confirmed to eliminate any false positive peaks with negative ChIP values. Coordinates of all GMEs are available in **Supplemental data 2**.

<u>GME associated gene analysis was performed as followed:</u> We first curated a list of the neuronal genes from the 17874 genes on Flybase. Information about Expression data, Gene Ontology terms and phenotypic information were batch-extracted from Flybase, resulting in 14824 genes with discernible information. We used a specific keyword list, including lncRNAs involved in neurogenesis, to further narrow the list to genes involved in developmental processes and embryogenesis (Stage 8 to Stage 17) (McCorkindale et al., 2019). The key word search list and final curated gene lists is available in **Supplemental data 1 and 3**.

### Hi-C

We adapted published Hi-C protocols (Tan et al., 2021; Zhang et al., 2020) to optimize for low- input. Around 15,000 FAC-sorted cells were used per replicate.

<u>Cell lysis:</u> Fixed cells were lysed in 600ul of ice-cold, freshly prepared Hi-C lysis buffer (0.2% Igepal, 10mM Tris pH 8.0, 10mM NaCl) with protease inhibitors (Sigma, 11873580001) on ice for 15 minutes. Cells were pelleted at 2500xg for 5min at 4°C and resuspended in Hi-C lysis buffer without protease inhibitors for a 2nd wash. Buffer was removed, leaving ∼10ml of solution. All subsequent centrifugation steps were performed at 4°C. Cells were incubated in 50ul of 0.5% SDS at 62°C for 10 min followed by SDS quenching with 1.14% TritonX-100 at 37°C for 15 min at 500rpm using a ThermoMixer.

<u>Chromatin digestion:</u> Next, 25ml of 10X NEBuffer DpnII and 10 ml of DpnII (50u/ml, NEB, R0543M) were added, and chromatin was digested overnight at 37°C, 500rpm. The enzymatic reaction was pelleted at 1000xg for 5 min. Supernatant was removed, leaving ∼27ml in the tube. <u>Biotinylation and ligation:</u> Digested ends were biotinylated with 0.4mM biotin-14-dATP (Fisher,19524-016), 10 mM dNTP mix (dC/G/TTP, NEB, N0446) and 0.8ml of 5U/µl DNA Polymerase I, Large (Klenow) Fragment (NEB, M0210) by incubating at 37°C for 1.5hrs at 300 rpm, shaking. Following biotinylation, 1ml of cold freshly prepared ligation buffer (1X T4 10X DNA ligase buffer, NEB B0202; 0.1 mg/ml BSA, NEB B9000) was added and reaction centrifuged at 1000xg for 5min. Supernatant removed, leaving ∼50ml. Ligation was performed in a 16°C water bath for 4hrs in 1ml of Ligation buffer supplemented with 10ml of 1U/ml T4 DNA ligase (Thermofisher, 15224-025). Tubes were occasionally inverted during incubation. Reaction was centrifuged for 5 min at 1000xg leaving ∼20ml of solution. Non-ligated ends were removed with Lambda Exonuclease (NEB, M0262) and Exonuclease-I (NEB, M0293) by incubating for 20 min at 80°C. At this step samples can be frozen at -20°C overnight.

<u>Tn5 transposition:</u> Cells were washed with Dip-C Wash buffer (25mM DTT, Sigma, 43816; 20mM Tris pH8.0, 20mM NaCl, 1mM EDTA) by centrifugation for 10min at 1000xg. ∼10ml of solution was left in the tube. Cells were lysed with 4ul of Dip-C Lysis buffer (25mM DTT, 20mM Tris pH8, 0.15% Triton X-100, 500nM Carrier ssDNA, 20mM NaCl, 1mM EDTA, 15ug/ml Protease, Qiagen, 19155) for 1hr at 50°C followed 70°C for 15 min (heat inactivation). The DNA was transposed by adding 15ml of 2X Transposase Buffer and 0.5ml of Tn5 Transposase (Diagenode, C01070012) to the reaction, incubated at 55°C for 10 min, and stopped with a 4ml of Stop Mix (45mM EDTA, 300mM NaCl, 0.01% Triton X-100, Protease 100mg/ml) by incubating for 40 min at 50°C (removal of Tn5), followed by incubation at 70°C for 20 min (protease inactivation).

<u>Streptavidin purification:</u> 25ul of Dynabeads MyOne Streptavidin T1 (Thermofisher, 65602) were washed with Tween Wash Buffer (TWB, 5mM Tris pH7.5, 0.5mM EDTA, 1M NaCl, 0.05% Tween- 20), resuspended in 34ml 2X Binding Buffer (10mM Tris pH7.5, 1mM EDTA, 2M NaCl) and added to the transposed DNA for biotin-streptavidin binding. DNA and beads were incubated at RT for 15 min with rotation. After 2X washes (ThermoMixer 55°C, 2min, 300rpm) with TWB, the beads were resuspended in 26.5ml of Nuclease-free water.

<u>Library preparation and sequencing:</u> DNA amplification was performed by PCR using indexed primers (Diagenode, C01011032). PCR reaction was separated on magnet, supernatant was purified using DNA clean &Concentrator -5 (Zymo Research, D4014). The purified library was further size-selected using AMPure XP beads at a 0.65X ratio, and the concentration and fragment size distribution were confirmed on the Bioanalyzer 2100. Libraries were pooled and sequenced pair-ended on NexSeq1000 Illumina Sequencer.

### Hi-C sequence alignment and processing

For each replicate, raw sequencing reads R1 and R2 were locally aligned to the dm6 reference genome independently using BWA (Li and Durbin, 2009) (*mem -A 1 -B 4 -E 50 -L 0*). The resulting SAM files were converted to BAM files using Samtools (Danecek et al., 2021). All the following steps were performed by HiCExplorer v3.7.2 (Ramirez et al., 2018). HiC matrices were generated by the *hicBuildMatrix* function at 25 kb binning (other options: *-seq GATC --danglingSequence GATC --minDistance 300 --maxLibraryInsertSize 1500 --minMappingQuality 10*). HiC matrices were filtered to only keep chromosome chr2L, chr2R, chr3L, chr3R, chrX, chr4 and chrY using the *hicAdjustMatrix* function. After analyzing the correlation between replicates (Correlation score, Hox cluster interaction and TAD borders overlap), HiC matrices from biological replicates were summed (merged replicates) using the *hicSumMatrices* function. Next, merged replicates were normalized between conditions for further analysis using the *hicNormalize* function ( *-n smallest*) and then corrected using the *hicCorrectMatrix* function (*correct*) performing the Knight-Ruiz matrix balancing algorithm (KR). To correct the effect of proximity ligation, the observed vs. expected values were computed (Obs/Exp) using the *hicTransform* function in HiCExplorer using the obs_exp method option. For biological replicates that were sequenced twice (technical replicates), reads were pooled at the raw sequencing reads steps (FASTQ files). Quality control metrics are available in **Supplemental data 4**.

### Hi-C data analysis

<u>Hi-C correlation:</u> Correlation between HiC biological replicates were assess using HiCREP (Yang et al., 2017). Briefly, HiC data at 25 kb resolution in cool format were imported into R using cool2matrix function and assess for correlation using the following HiCREP parameters: h=10, lbr=3 x resolution (75000) and ubr= 10 x resolution (250000). Correlation scores “scc” were displayed as a matrix using (Galili et al., 2018).

<u>Average GME interaction:</u> Average contact interactions between all detected GMEs were performed using the R package GENOVA (van der Weide et al., 2021). Briefly, KR or OE Hi-C matrices, resulting from the above HiCExplorer pipeline, were converted into COOL format using the *hicConvertFormat* function *(--inputFormat h5 --outputFormat cool*) and imported into GENOVA using the *load_contacts* function, with the balancing option at *true* or *false,* respectively. Chromosome sizes from *dm6 Drosophila* reference genome, centromere regions, and the list of regions of interest (GME, PRE or random genomic regions) were imported into GENOVA as well. Bin indices were synchronized between conditions using the *synch_indices* function. All-vs-all GME-GME contacts were computed and displayed as an average submatrix using the *PESCAn* function without threshold of distance for OE matrices input or excluding any interaction below 300 kb of distance for KR corrected Hi-C matrices. The shifted backgrounds, which computes the average Hi-C signal at a fixed distance from all individual GME-GME interactions, were computed at 100 kb, 500 kb and 1Mb from GME contact site. To filter for *cis* (intra chromosomal arm interaction) or *trans* interactions (inter chromosomal arms and inter chromones) the variable *mode* from the *anchors*_*PESCAn* function was modified accordingly. Averaged interaction submatrices for signal and background were displayed using ggplot2 (Villanueva, 2019).

<u>GME contact detection:</u> Detection of significant contacts was performed by FitHiC2 (Ay et al., 2014; Kaul et al., 2020), transforming each Hi-C bin values into a probability interaction based on the level of the expected interaction. Due to the limited number of starting material from FAC- sorted neuroblasts, HiC-ACT was used to refine the probability of contact detection based on local probability distribution of each bins (Lagler et al., 2021). Briefly, normalized un-corrected .h5 Hi- C matrices from HiCexplorer were transformed into .cool format data using *hicConvertFormat* function and then converted into a .bedpe format ({chr1,start1,end1,chr2,start2,end2,value} using the *dump* function from Cooler (dump --join) (Abdennur and Mirny, 2020).

The data were further reduced into a {chr1,start1,0,chr2,start2,0,value} format using awk. After a by-chromosome sort (*sort -k2,2d -k6,6d*), Juicer tools *pre* function (Durand et al., 2016) was used generating a Juicer HiC matrix .hic file format ready to be imported into FitHiC2 using the built-in *createFitHiCContacts-hic.py* script that were contacted on all possible chromosomal interactions between chrX, 2L, 2R, 3L and 3R *(--resolution 25000 --datatype observed --Norm NONE*). Next, the imported Hi-C data were KR corrected using the HiCKRy script generating the biases (https://github.com/ay-lab/HiCKRy).

Lastly, Hi-C fragment files at 25 kb resolution were generated using the *createFitHiCFragments- fixedsize.py* function. The corrected Hi-C data, biases and fragment files were input into fithic.py script with the following options: *--lowerbound 50000*, contacts below 50 kb are not considered, *- x all* which computed contact for intra and inter chromosomal interactions. FitHiC2 result was then refined using HiC-ACT with the following options and parameters: *kb=25000* (resolution), *h=11* (recommended smoothing for 25 kb resolution), threshold=0.05 (*P* value for FitHiC2 result filtering). The resulting refined HiC-ACT matrices were thresholded for *P* value<0.01 to generate the list of significant contacts. Significant contacts submatrices were visualized individually or as an Aggregate Peak Analysis using the APA function of GENOVA.

<u>TAD detection:</u> Detection of TAD borders and TAD domains were performed using the *hicFindTADs* function with the following options: --correctForMultipleTesting fdr --minDepth 75000 --maxDepth 2500000 --step 25000. TAD border regions have the size of the resolution (25 kb) whereas TAD domain regions start and end at the middle of each flanking borders. TAD borders were detected for each neuroblast Hi-C replicates and overlap analysis was performed using *intervene venn* tools with all default options. To analyze the number of TADs that each of the FitHiC2::HiC-ACT significant contacts were spanning, each significant contact was assigned to its encompassing TADs (TAD’s ID were numbered sequentially along each chromosomal arms) and each significant contact was assigned a *inter* or *intra-TADs* label.

<u>GME clustering analysis:</u> To calculate the proportion of GMEs that interact with each GME, a virtual 4Cs of all GME were extracted from the Hi-C data. For GME clustering analysis, only contacts involving GMEs were studied. The 4C of each GME viewpoint was compared to the 4C of each of its contacts, and the proportion of shared direct contacts were quantified. We performed the same operation for all GMEs that did not contact that viewpoint GME. Then the two datasets were compared to quantify the degree of overlap in the GME 4C network between contacting and non-contacting GME pairs.

### Other Statistical analysis

We applied standard t-tests (with var.equal = TRUE) and Wilcoxon Rank Sum and Signed Rank test (default parameters) on R. Statistical significance was classified as follows: ^∗^ < 0.05, ^∗∗^ < 0.01, ^∗∗∗^ < 0.001, ^∗∗∗∗^ < 0.0001. For ParS experiment statistical tests, each data point represents an embryo, and the number of ParS signals quantified is shown on the graph. Random sampling (bootstrap simulation) was performed on R and the estimate the P value of bootstrap is *P value = (1 + sum(s >= s0)) / (N+1)*, in which *s* is random values and *s0* is the observed value.

### Lamin:DamID

<u>Embryo collections:</u> Two populations cages were prepared, one containing UAS-Dam only flies (control) and the other containing UAS-Dam::Lamin fusion flies. Both were crossed to Wor-GAL4 flies to express Dam or Dam::Lamin in neuroblasts. After a one-hour prelay, embryos were collected after one hour and aged according to desired stage. Early-stage DamID, embryos were aged at 29C for 4.5hours (st10 embryos-we age for 3hours), weighed, and frozen at -20C. For late stage DamID, embryos were first aged at 18C for 17.5hours and then incubated at 20C for 4.5hours. For both, embryos were hand-cleaned during incubation to remove all dead embryos or embryos at the wrong stage.

<u>DamID processing:</u> Genomic DNA was processed following the protocols detailed in (Sen et al., 2019; Southall et al., 2013). Briefly, genomic DNA was extracted using a Qiagen kit, digested with DpnI (NEB Cat# R0176) and purified (Qiagen PCR purification Kit #28016). Digested fragments were ligated to adaptors with T4 ligase (NEB Cat# M0202M) overnight and then digested with DpnII (NEB Cat# R0543S). Methylated fragments were PCR amplified with MyTag, PCR cleaned, and sonicated (2ug of purified DNA in CutSmart buffer was sonicated with Covaris peak power 140, Duty Factor 10, 200 cycles, average time 130sec). Sonicated products were checked with a Bioanalyzer and confirmed to be ∼300bp size. These fragments were used for library preparation and sequencing.

### Confocal imaging and image analyses

Zeiss 700 Axio Imager 2 laser scanning confocal was used for embryo imaging. ParB/ParS images were taken at a 0.4µm step size, and pinholes were adjusted to have equal optical section thickness for all channels. All image analyses were done on Fiji (Schindelin et al., 2012).

<u>H3K27me3 quantification:</u> Quantification of H3K27me3 signal in nuclei of neuroblast or epithelial cells was perform using TrackMate tools in Fiji (Ershov et al., 2022). Briefly, the 3D imaging data of the H3K27me3 channel was extracted and the outside of the nucleus was set to 0 so that only signals from the nucleus were used for quantification. TrackMate works with space and time data to segment objects and analyze trajectory. The z-axis parameter was switched to T (time). The LoG algorithm was used to segment the H3K27me3 foci with an approximated radius of 0.5µm and the median filter was set to 1. To link the detected foci through Z planes, the Simple LAP algorithm was used to track foci using the following paramters: Linking = 0.11, Gap = 0 and Frame Gap = 0. TrackMate results were analyzed on R. Between 15 and 18 cells were analyzed per embryo for a total of 28 neuroblasts and 30 epithelial cells.

*<u>ParB/Lamin proximity quantification:</u>* After selecting each neuroblast from the ventral nerve cord that are in interphase (excluding neuroblast in mitosis undergoing nuclear envelope breakdown based on Lamin signal and nucleus size) we identified the z plane in which the ParB foci was strongest and measured the shortest distance to the nuclear lamina. ParB foci for which pixels overlapped with those of Lamin signal were scored to be “on” Lamin, and ParB foci not touching Lamin but still within 0.4µm of the lamina were scored as “near.” The proportion of ParB foci at the lamina was calculated as being the summed of the score of “on” and “near” divided by the total number of foci. Around 50 neuroblasts were analyzed per embryo with at least 3 embryos per condition.

<u>GME-insertion GFP quantification :</u> Quantification of the plasma membrane anchored mCD4::tdGFP from *dpn-{IVS/GME}-mCD4::tdGFP* transgenes was performed by identifying a z plane in which most of the neuroblasts were visible (by Worniu staining). The GFP signal was quantified on the GFP channel by measuring the average intensity in a 4x4 pixel square at 102 random locations at the plasmid membrane of various neuroblast spanning the whole ventral nerve cord. The same process was performed to evaluate the GFP background by measuring the intensity of GFP at 50 random locations across the embryo outside of the cells. The background corrected GFP intensities were used for statistical test. To visually help the detection of the GFP an inversed enhanced GFP channel was generated. Three replicates were quantified per condition.

### ChIP-seq analysis

Raw ChIP-seq datasets from Brown et, al. 2018 (GSE102339) (Brown et al., 2018) for E(z), Pc, Ph, Psc, Pho and input and Bonnet et, al. 2019 (GSE114832) (Bonnet et al., 2019) for Pho and input were processed as follows: adaptor removal using Cutadapt (-m 5, -e 0.2), alignment to dm6 using bowtie2 (-local, --very-sensitive-local), removal of PCR duplicates using PICARD Mark duplicates, removal of mitochondrial DNA and read quality filter at 30 using Samtools (-q 30). Resulting bam files were merged by replicate, and log2 (RPM-normalized ChIP) / (RPM- normalized Input) bigwig files were generated using Bedtools (bamcompare). We identified putative Pho motifs occurrence within the *hb* genomic region using PWMScan (PWMTools, ccg.epfl.ch) (Ambrosini et al., 2018) by selecting Pho_DROME_B1H OTF0277.1 (OnTheFly 2014 motif library) with a p-value cutoff at 5.10^-4^.

## Supplementary figures

**Supplemental Figure 1.**
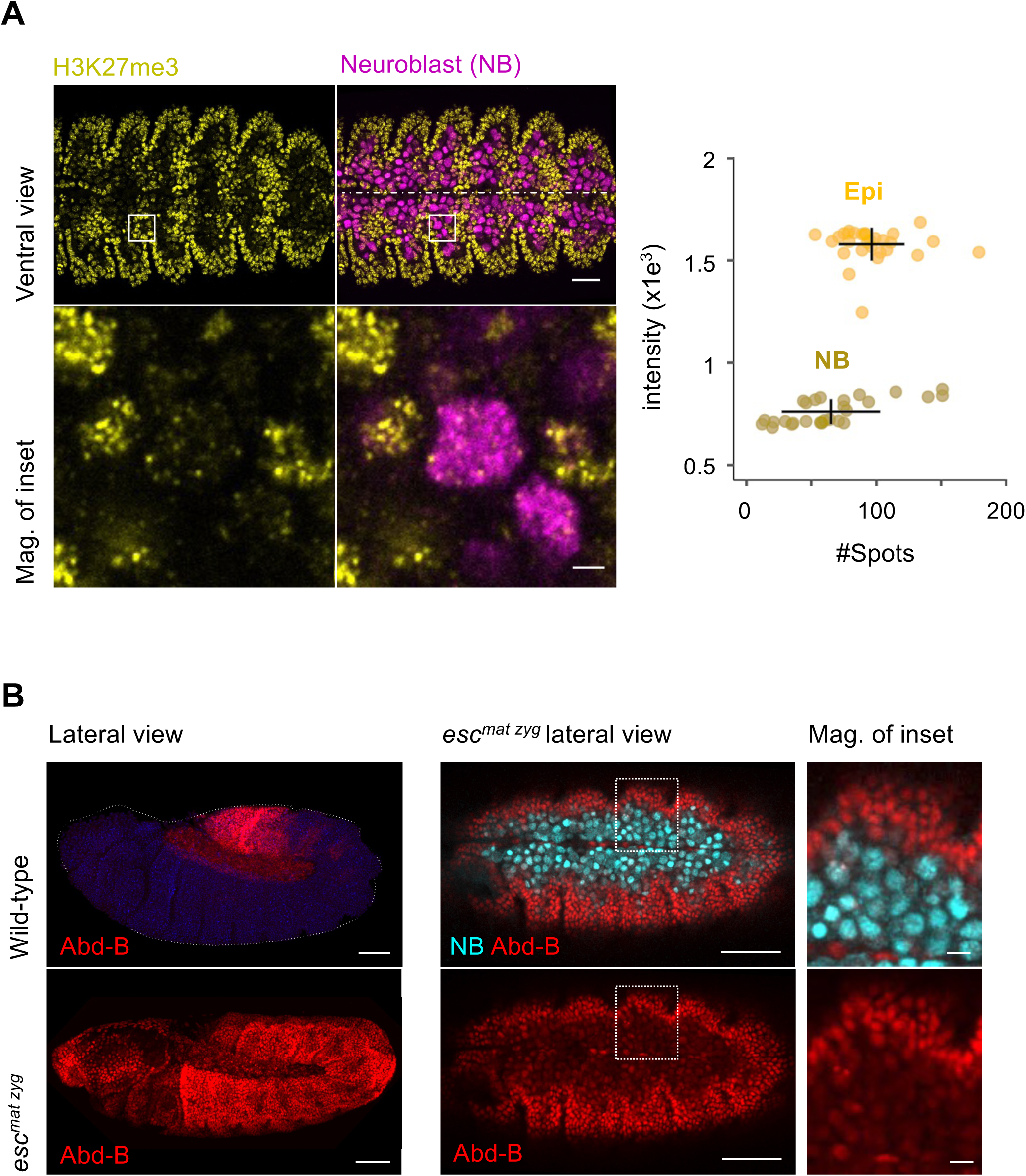
(Supplemental to Figure 1): **A. H3K27me3 signal is decreased in neuroblast:** *Left*, Immunostaining of the Polycomb repressive complex 2 epigenetic mark H3K27me3 in stage 14 embryos. Ventral single Z-plane showing pan-neuroblast Worniu marker (Wor) in magenta and the H3K27me3 marks in yellow. The Embryonic midline is shown as a dashed white line. Scale bar = 20µm, scale bar of close up = 2µm. *Right*, scatter plot quantification of H3K27me3 foci in neuroblast vs non neuroblast showing the background normalized intensity of H3K27me3 staining signal in function of the number of H3K27me3 detected foci (#Spots). Standard deviations are shown as vertical and horizontal black lines and average at the intersection. **B. Lack of Abd-B derepression in PRC2 mutant embryo:** Immunostaining of wild-type and maternal and zygotic mutant of the PRC2 subunit extra-sex-combs, *esc*. *Left*, Side views of wild-type and *esc* mutant embryos stained for Abd-B in red, and DAPI in blue (wild-type only). Scale bars = 50µm. *Middle*, Single Z-plan of ventral view of the *esc* mutant embryos stained for Abd-B in red and pan-neuroblast marker Wor in cyan. Scale bar = 50µm. *Right*, magnification of the insets from middle panels, Scale bar = 20µm.

**Supplemental Figure 2.**
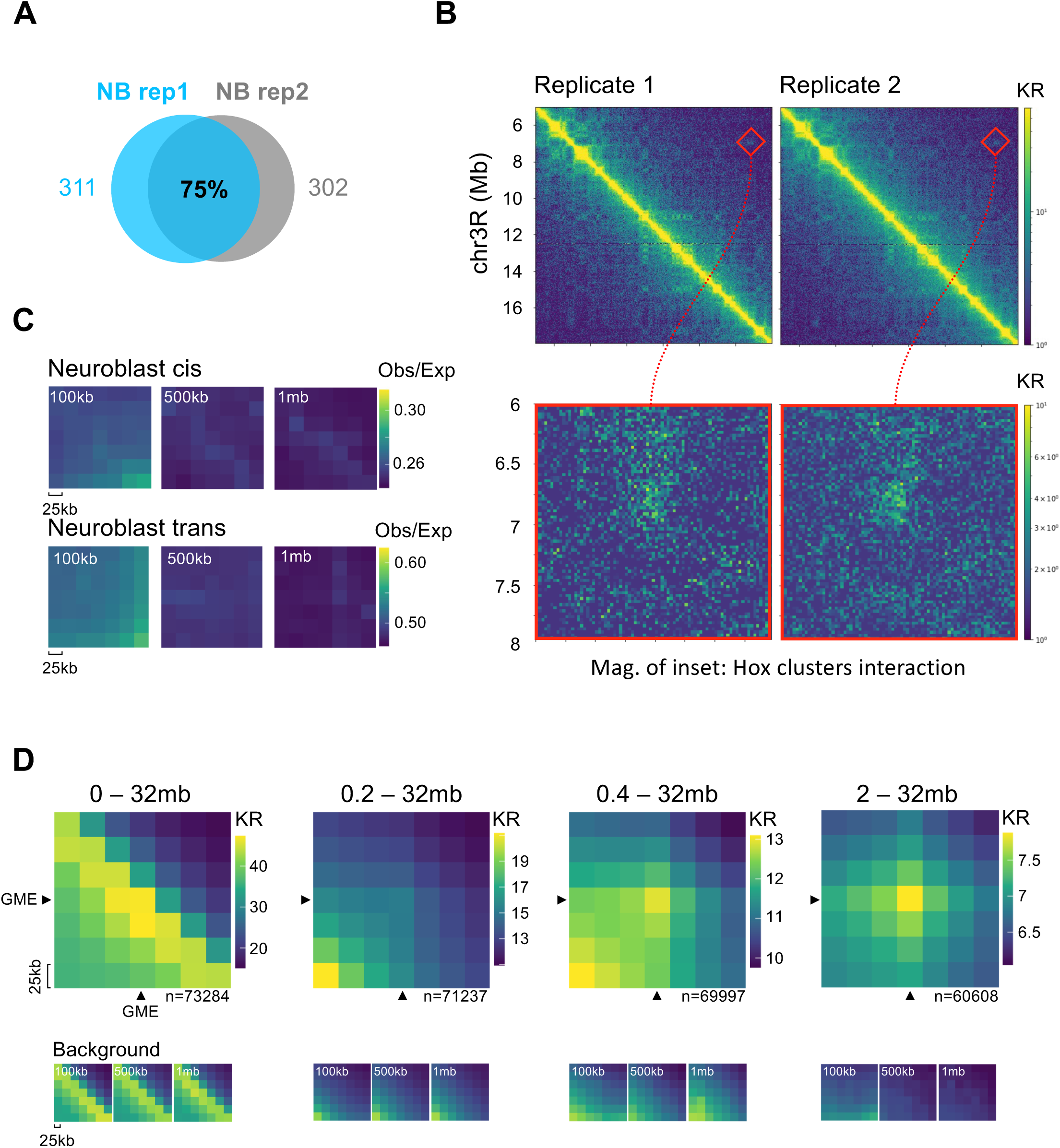
(Supplemental to Figure 1): **A. Strong TAD borders overlap among Hi-C neuroblast replicates:** Venn diagram showing the overlap of detected TAD borders in neuroblast from two Hi-C biological replicates 25 kb resolution of embryonic neuroblast stage 10. Hi-C maps were normalized and KR-corrected. **B. Hox clusters interactions are detected in neuroblast Hi-C replicates:** *Top*, 25 kb resolution Hi-C matrices from two biological replicates of neuroblast embryonic stage 10 spanning ANT-C and BX-C Hox gene clusters on chromosome 3R (highlighted by the red squares). *Bottom*, magnification of Hox cluster interaction. Hi-C maps were normalized and KR-corrected and shown as log1p scale. **C. Background at GME-GME interaction:** Average submatrices, sampled at distances of 100 kb, 500 kb, and 1 Mb from the GME-GME cis or trans interaction in neuroblast. 25 kb resolution Hi-C matrices were KR-correct and the Observed/Expected (O/E) is shown. **D. GME-GME cis interactions are detected in neuroblasts :** Average submatrices of all possible GME-GME cis interactions in neuroblasts at various distance ranges. GME- containing Hi-C bins are pointed by black arrows. The total number of GME-GME interactions (n=) are shown and the background average submatrices, sampled at distances of 100 kb, 500 kb, and 1 Mb from GME-GME interactions are shown below each GME-GME average Hi-C submatrices. 25 kb resolution Hi-C matrices were KR-correct.

**Supplement Figure 3.**
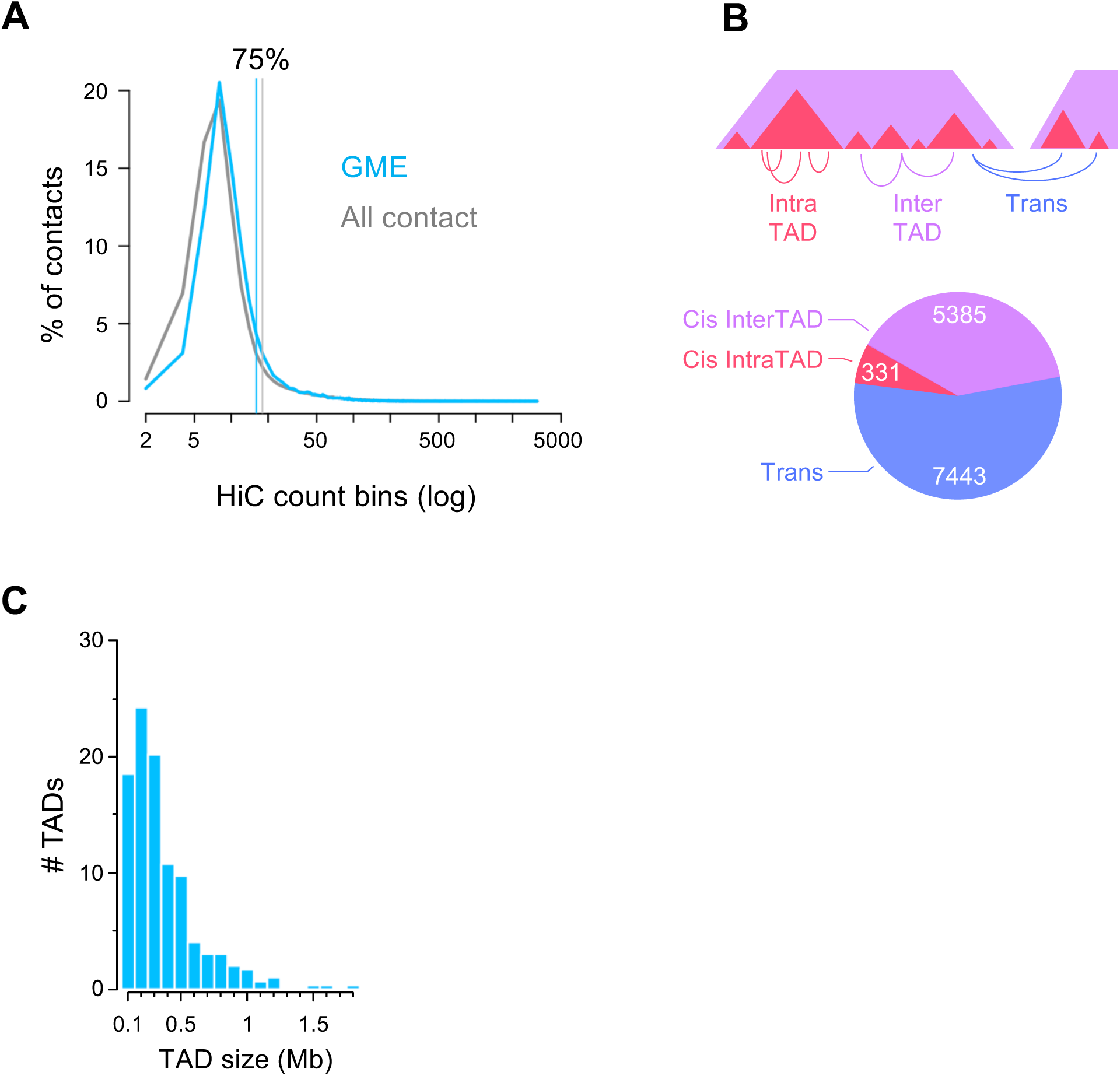
(Supplemental to Figure 2): **A. GME significant contacts and all significant contacts Hi-C count:** Distribution of the proportion of significant contacts in function of Hi-C count (KR-corrected) shown on a log scale. Horizontal lines mark the 75th percentile of the distributions. **B. GME-GME contact detection:** *Top,* Schematic representation of contact types detected in the Hi-C data. Cis contacts connect regions within the same chromosome: cis, intra contacts occur within the same TAD, while cis, inter contacts span across different TADs on the same chromosome arm. Trans contacts connect regions from different chromosomes or chromosome arms. *Bottom*, Circular diagram showing the number of significant contacts per type. **C. TAD size distribution:** Distribution of the size of detected TAD (n=297) genome wide (mean ∼400 kb, median of ∼350 kb).

**Supplemental Figure 4:**
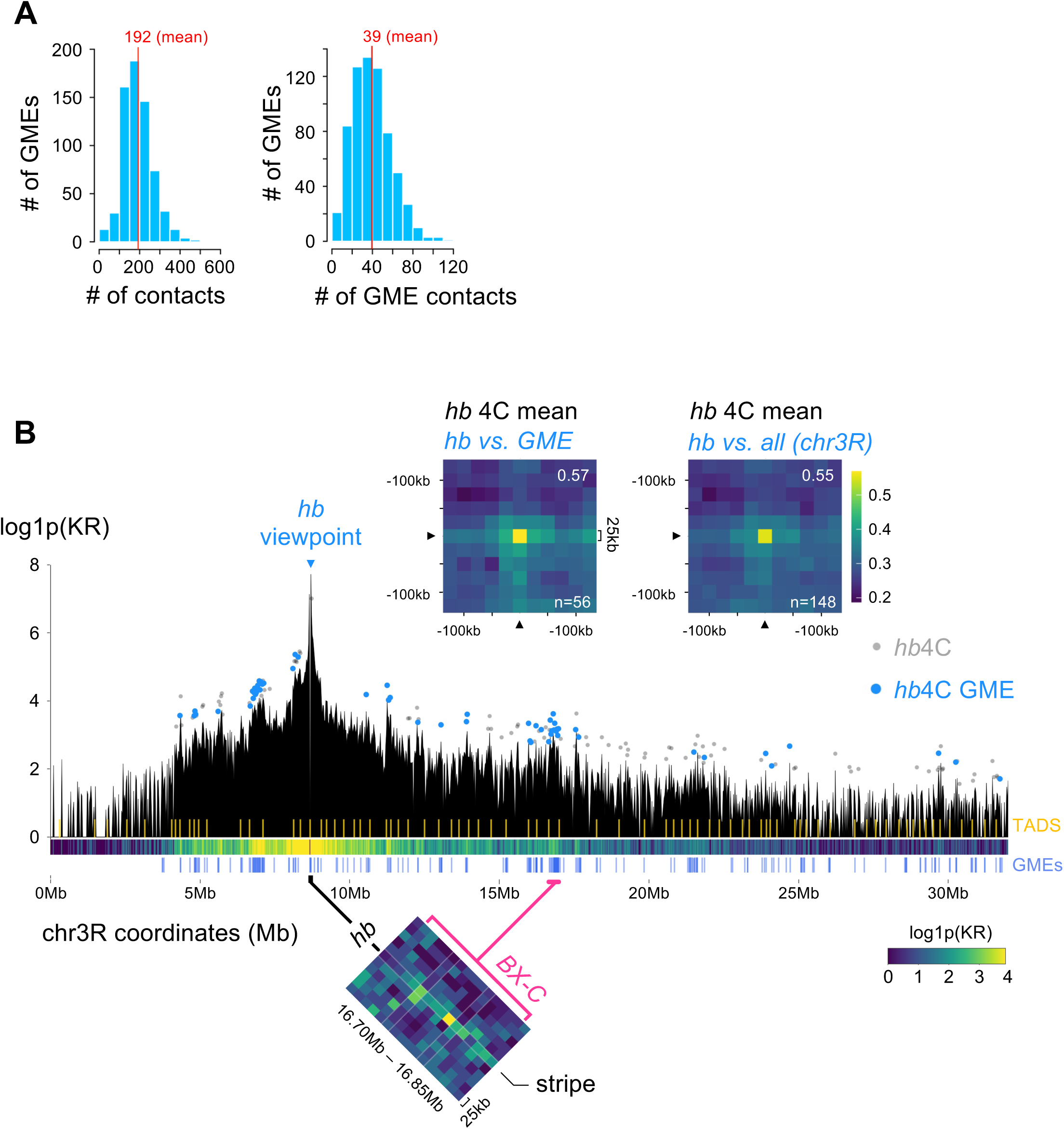
(Supplemental to Figure 2): **A. GME Contact Statistics:** *Left*, Distribution of significant contacts from the virtual 4C of each GME viewpoint. *Right*, Distribution of significant GME-GME contacts within the virtual 4C of each GME viewpoint. **B. *hunchback* 4C:** Virtual 4C analysis of the *hb* GME on chromosome 3R from KR-correct neuroblasts Hi-C matrix at 25 kb resolution. Contact types: Contacts involving GMEs are shown in blue, while non-GME contacts are depicted in grey. Yellow lines indicate Topologically Associating Domain (TAD) borders. GMEs positions are shown as blue vertical bars. *Top,* Average Hi-C submatrices of all the 4C contacts involving the *hb* GME and either other GMEs (*top*-*left*) or all other contacts (*top*-*right*) . Hi-C bins involved in the significant contacts are pointed by black arrows. Total number of contacts are shown in white. *Bottom,* Zoom-in of the contact between the GME of *hb* and the BX-C Hox cluster GMEs. KR-corrected and Observed/Expected values are shown.

**Supplemental Figure 5.**
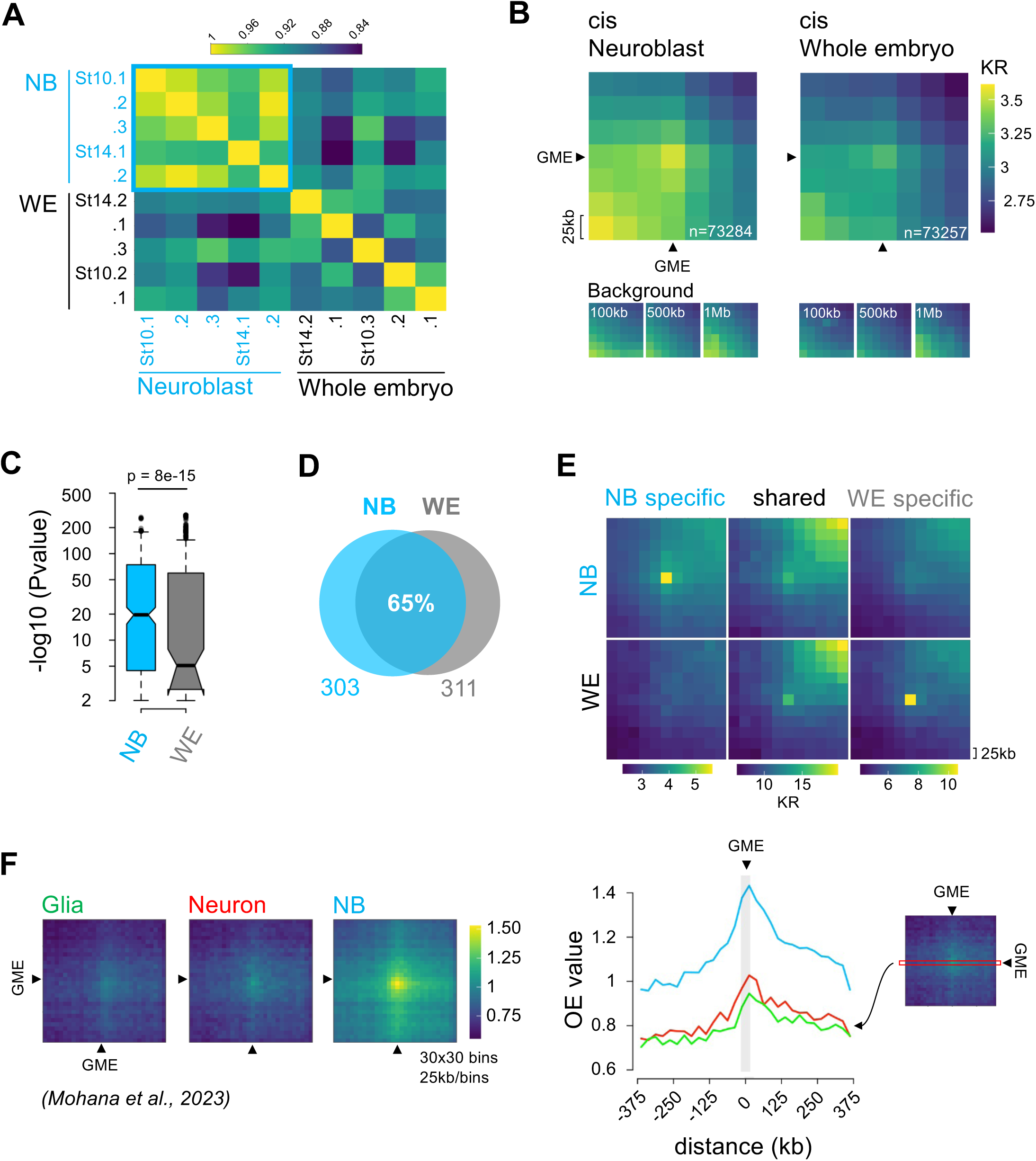
(Supplemental to Figure 3): **A. Hi-C sample correlation:** Correlation heatmap showing HiCREP *scc* scores for neuroblast (blue labels) and whole embryo (black labels) Hi-C replicates at 25 kb resolution. Hi-C data were unnormalized and uncorrected. **B. GME-GME interactions are stronger in neuroblast than whole embryo:** *Top*: Average Hi-C submatrices of all possible GME-GME cis interactions in neuroblast and whole embryo cells. 25 kb resolution HiC matrices were normalized and KR-corrected. GME-containing Hi-C bins are pointed by black arrows. Total number of GME-GME interactions are shown in white. *Bottom:* Average submatrices, sampled at distances of 100 kb, 500 kb, and 1 Mb from the GME-GME interaction. **C. GME-GME contacts are more significant in neuroblast than whole embryo:** Distribution of significant cis contacts p-values in neuroblast and whole embryo. Wilcoxon test was used for significance. **D. TAD borders overlap:** Venn diagram showing the overlap of detected TAD borders in neuroblast and whole embryo at 25 kb Hi-C resolution. Hi-C maps were normalized and KR-corrected. **E. Neuroblasts make cell-type specific GME-GME contacts:** Average Hi-C submatrice of neuroblast specific, whole embryo specific and shared significant GME-GME contacts in neuroblast and whole embryo. Hi-C data were normalized and KR-corrected values were displayed. Analyzed contacts were thresholded by distances ranging from 0.4 Mb to 32 Mb. **F. GME-GME contacts are stronger in neuroblast compared to other CNS cell types:** Average Hi-C submatrice of neuroblast GME-GME significant contacts in glia, neurons and neuroblast from Mohana et al., 2023 (GSE214707). 25 kb resolution Hi-C data were normalized, KR-corrected and observed/expected values were displayed.

**Supplemental Figure 6.**
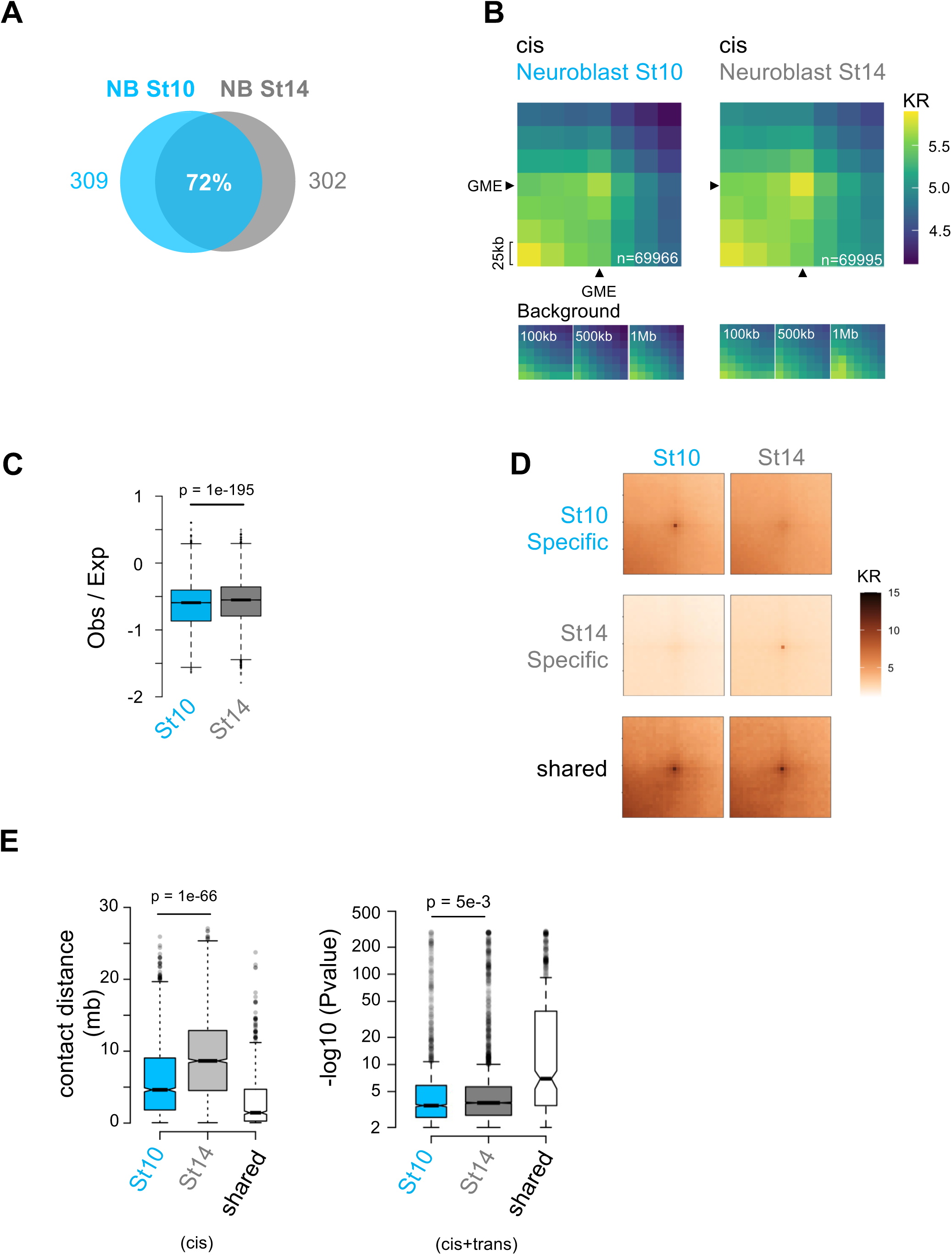
(Supplemental to Figure 4): **A. Neuroblasts TAD border overlap:** Venn diagram showing the overlap of detected TAD borders in neuroblast stage 10 and14 at 25 kb Hi-C resolution. Hi-C maps were normalized and KR-corrected. **B. Cis GME-GME interaction over time:** *Top*: Average Hi-C submatrices of all possible GME- GME cis interactions in neuroblast stage 10 and 14. 25 kb resolution HiC matrices were normalized and KR-corrected values were displayed. GME-containing Hi-C bins are pointed by black arrows. Total number of GME-GME interactions are shown in white. *Bottom:* Average submatrices, sampled at distances of 100 kb, 500 kb, and 1 Mb from the GME-GME interaction. **C. Strength of *cis* GME-GME interaction:** Boxplot comparing the distribution of Hi-C Observed/Expected values of neuroblast stage 10 and stage 14. Stage 10: n=73253 interactions, including 12428 null. Stage 14: n=73282 interactions, including 8108 null interactions. Wilcoxon test was used for significance. **D. Stage specific GME contacts:** Average Hi-C submatrice of neuroblast stage 10 specific, stage 14 specific and shared significant GME-GME contacts in neuroblast stage 10 and 14. Hi-C data were normalized and KR-corrected values were displayed. Analyzed contacts were thresholded by distance ranging from 0.4 Mb to 32 Mb. **E. Stage specific and shared GME-GME contact distance distribution and P-value:** Boxplots comparing the distance (left) or the Pvalue (right) distribution of cis stage-specific and stage-shared GME-GME contacts. Wilcoxon test was used for significance.

**Supplemental Figure 7.**
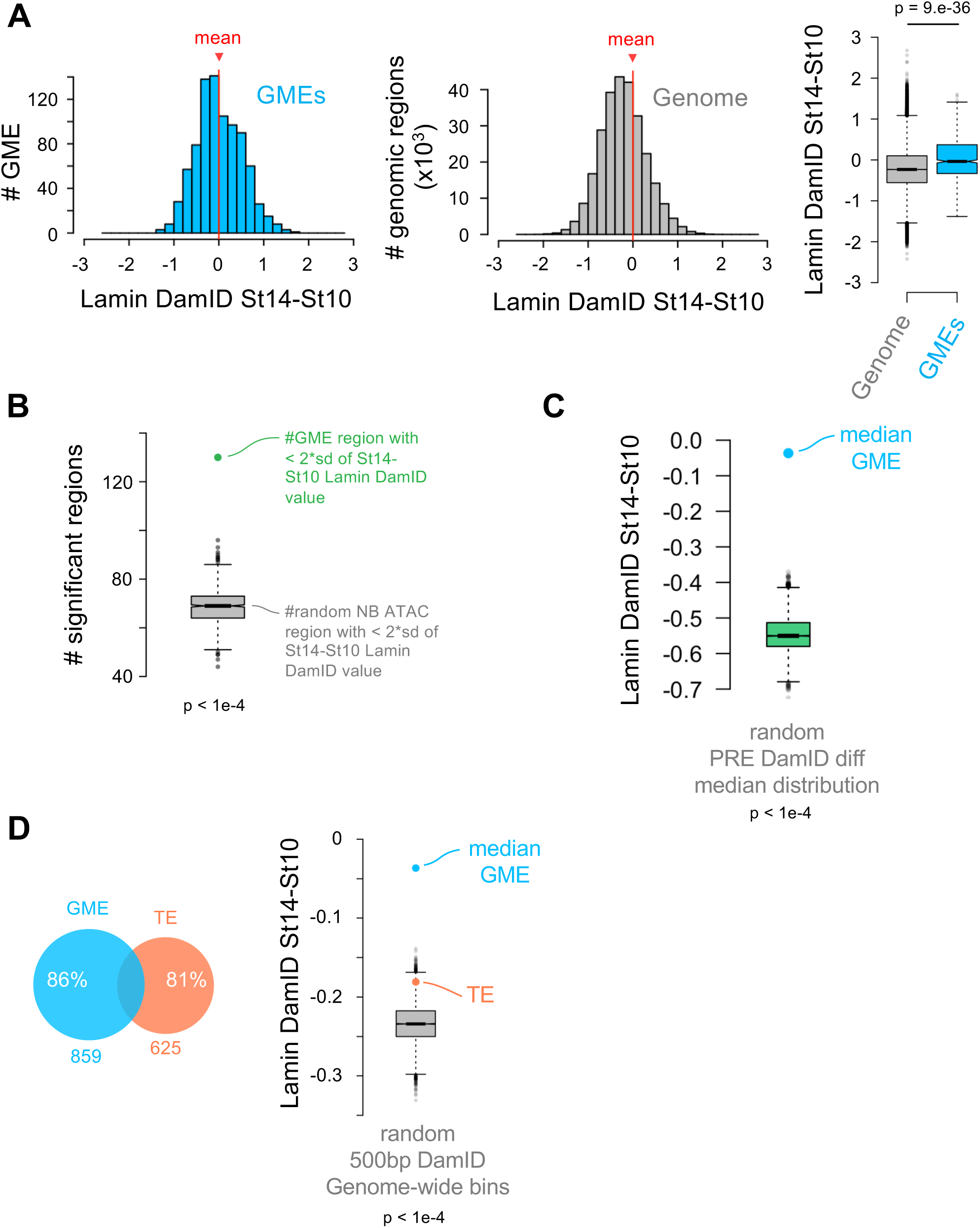
(Supplemental to Figure 4): **A. Lamin DamID distribution at GMEs and genome-wide:** Distribution of GMEs (Left) or whole genome (Middle) relative to their differential Lamin DamID signal (DamID stage 14 - stage 10); Means are displayed as red vertical lines and arrows. Right: Boxplot showing the distribution of the differential Lamin DamID signal genome-wide compared to GME regions. Wilcoxon test was used for significance. **B. Lamin DamID at GME vs non GME:** Boxplot comparing the number of GME for which the differential DamID value is exceeding 2 standard deviations above the genome-wide average, green dot (see figure 4E) to the distribution of median of differential DamID value is exceeding 2 standard deviations above the genome-wide average from randomly sampled ATAC open sites excluding GME regions. 10000 random sampling of 859 ATAC open site were performed (bootstrap p-value < 1e-4). The black bar represents the median of the random distribution. **C. Lamin DamID at GME vs PREs:** Boxplot comparing the median of the differential DamID of GMEs to that of the distribution of median from randomly sampled PREs. 10000 random sampling of 859 PREs were performed (bootstrap p-value < 1e-4). The black bar represents the median of the random distribution. **D. GME versus TE comparison**: *Left*: Venn diagrams showing the overlap between GME and Tethering Elements (TE) from Batut et al., 2022. *Right*: Boxplot comparing the medians of the differential DamID of GMEs and TEs to that of the distribution of median from randomly sampled genome-wide 500bp regions. 10000 random sampling of 625 genome-wide regions were performed (bootstrap p-value < 1e-4). The black bar represents the median of the random distribution.

**Supplemental Figure 8.**
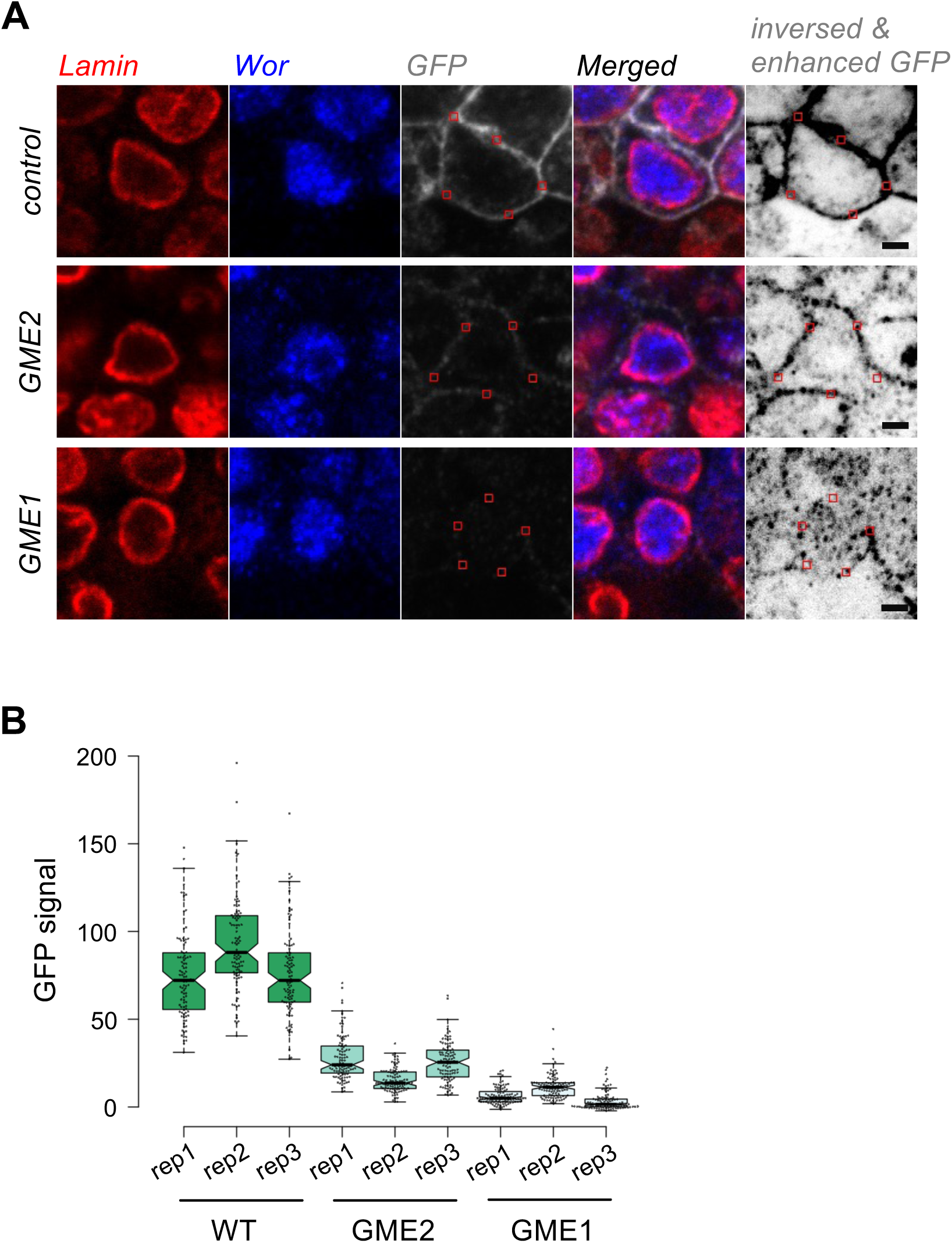
(Supplemental to Figure 5): **A. Example of GFP quantification from mCD4 transgenes:** Representative images of neuroblasts used for mCD4-GFP quantification, including Lamin (red), Worniu (blue), GFP (circled in red), a merged image, and an inverted merged image with enhanced GFP signal. The outlined workflow highlights steps for isolating GFP signals for analysis. **B. mCD4 transgene expression quantification across embryos:** GFP signal intensity quantified on an 8-bit scale (0–255). For each embryo, GFP fluorescence from the dpn-mCD4- GFP transgene was sampled at 100 random positions along the neuroblast plasma membrane and at 50 random background positions (outside cells). Plasma membrane intensity was background-corrected by subtracting the average background signal. Scale bar: 2 μm.

