## Supplemental_data_1 for "Gene mobility elements mediate cell type specific genome organization and radial gene movement *in vivo*"

**Keywords for neuronal gene list curation:**

Head | head | Neuron | neuron | Neurons | neurons | Neuroblast | neuroblast | neural | Neural | Ganglion | ganglion | Cord | cord | Interneuron | interneuron | Motor neuron | motor neuron | Sensory neuron | sensory neuron | Glia | glia | Glial | glial | Sensory | sensory | Nervous | nervous | Ectoderm | ectoderm | Brain | brain | Nerve | nerve | Cephalic | cephalic | Procephalic | procephalic | Neuronal | neuronal | CNS | PNS | Axon | axon | Axonal | axonal

**Keywords for neuronal developmental gene list curation:**

Development | development | Develop | develop | Segmentation | segmentation| Segment |segment | Specification | specification | Generation | generation | Generate | generate| Generates | generates | Patterning | patterning | Fate determination | fate determination | Determination | determination | Neural precursor | neural precursor | Neuroblast | neuroblast | Primordium | primordium

**Keywords for neuronal developmental embryonic Stage 8 to 16 gene list curation:**

embryonic stage 8 | embryonic stage 9 |embryonic stage 10 | embryonic stage 11 | embryonic stage 12 | embryonic stage 13 |embryonic stage 14 | embryonic stage 15 | embryonic stage 16 | embryonic stage 14
